# Infection and transmission of ancestral SARS-CoV-2 and its alpha variant in pregnant white-tailed deer

**DOI:** 10.1101/2021.08.15.456341

**Authors:** Konner Cool, Natasha N. Gaudreault, Igor Morozov, Jessie D. Trujillo, David A. Meekins, Chester McDowell, Mariano Carossino, Dashzeveg Bold, Taeyong Kwon, Velmurugan Balaraman, Daniel W. Madden, Bianca Libanori Artiaga, Roman M. Pogranichniy, Gleyder Roman Sosa, Jamie Henningson, William C. Wilson, Udeni B. R. Balasuriya, Adolfo García-Sastre, Juergen A. Richt

**Author notes:** Institut für Virologie, Justus-Liebig-Universität, Giessen, Germany (Gleyder Roman Sosa). **Corresponding Author:** Juergen A. Richt, DVM, PhD, Department of Diagnostic Medicine/Pathobiology, College of Veterinary Medicine, Kansas State University, Manhattan, KS, USA.

## Abstract

SARS-CoV-2, a novel *Betacoronavirus*, was first reported circulating in human populations in December 2019 and has since become a global pandemic. Recent history involving SARS-like coronavirus outbreaks (SARS-CoV and MERS-CoV) have demonstrated the significant role of intermediate and reservoir hosts in viral maintenance and transmission cycles. Evidence of SARS-CoV-2 natural infection and experimental infections of a wide variety of animal species has been demonstrated, and *in silico* and *in vitro* studies have indicated that deer are susceptible to SARS-CoV-2 infection. White-tailed deer (*Odocoileus virginianus*) are amongst the most abundant, densely populated, and geographically widespread wild ruminant species in the United States. Human interaction with white-tailed deer has resulted in the occurrence of disease in human populations in the past. Recently, white-tailed deer fawns were shown to be susceptible to SARS-CoV-2. In the present study, we investigated the susceptibility and transmission of SARS-CoV-2 in adult white-tailed deer. In addition, we examined the competition of two SARS-CoV-2 isolates, representatives of the ancestral lineage A (SARS-CoV-2/human/USA/WA1/2020) and the alpha variant of concern (VOC) B.1.1.7 (SARS-CoV-2/human/USA/CA_CDC_5574/2020), through co-infection of white-tailed deer. Next-generation sequencing was used to determine the presence and transmission of each strain in the co-infected and contact sentinel animals. Our results demonstrate that adult white-tailed deer are highly susceptible to SARS-CoV-2 infection and can transmit the virus through direct contact as well as vertically from doe to fetus. Additionally, we determined that the alpha VOC B.1.1.7 isolate of SARS-CoV-2 outcompetes the ancestral lineage A isolate in white-tailed deer, as demonstrated by the genome of the virus shed from nasal and oral cavities from principal infected and contact animals, and from virus present in tissues of principal infected deer, fetuses and contact animals.

## Introduction

The family *Coronaviridae* is comprised of enveloped, single-stranded, positive-sense RNA viruses, and include four genera *Alpha-, Beta- Gamma- and Delta-coronaviruses. Betacoronaviruses* have been the subject of intensive research since the emergence of severe acute respiratory syndrome coronavirus (SARS-CoV) in 2002, Middle East respiratory syndrome coronavirus (MERS-CoV) in 2012, and most recently SARS-CoV-2 in 2019. In order to determine the origins of SARS-CoV-2, surveillance efforts have mainly focused on bat populations since they were identified as the reservoir species for SARS-CoV-like and MERS-CoV-like viruses^1^. Intermediate hosts such as civet cats for SARS-CoV or camels for MERS-CoV have also been identified as an important vehicle for virus spillover into human populations and have shown to play a significant role in pathogen establishment and continued animal-to-human transmission^2,3^.

The World Organization for Animal Health (OIE) has reported the natural infection of SARS-CoV-2 in at least 10 animal species across continents including the Americas, Europe, Africa and Asia: domestic cats and dogs, tigers, lions, cougar, snow leopard, puma, mink, ferrets, gorilla, and otter ((https://www.aphis.usda.gov/aphis/dashboards/tableau/sars-dashboard). In the United States alone, USDA-APHIS has reported 217 incidences of natural SARS-CoV-2 infections amongst 9 different species (www.aphis.usda.gov). Experimental infection of SARS-CoV-2 in animal models has identified cats, ferrets, mink, Syrian golden hamsters, non-human primates, tree shrews, and deer mice as highly susceptible to SARS-CoV-2 infection^4^. Dogs, cattle, and Egyptian fruit bats have shown moderate susceptibility while non-transgenic mice (with the exception of variants containing the N501Y polymorphism in their S gene), poultry, and pigs are not readily susceptible to SARS-CoV-2 infection^4^. It is important to determine susceptible host species for SARS-CoV-2 in order to better understand the ecology of this virus and to identify potential reservoir species which may be sources of spillover into human populations. Additionally, the emergence and sustained transmission of SARS-CoV-2 variants of concern (VOC) has important implications in virus evolution and pathogenesis^5^. It is therefore necessary to investigate the transmission efficiency and pathogenesis of SARS-CoV-2 VOCs in susceptible species.

A recent publication by Palmer and coworkers^6^ describes susceptibility of white-tailed deer (*Odocoileus virginianus*) fawns to the SARS-CoV-2 tiger isolate TGR/NY/20^6^. The work presented here expands upon the previous findings by describing SARS-CoV-2 infection in pregnant adult deer, as well as horizontal and vertical transmission of the virus. Furthermore, we investigated the competition of two SARS-CoV-2 isolates in deer, representatives of the ancestral lineage A (SARS-CoV-2/human/USA/WA1/2020) and the alpha VOC B.1.1.7 (SARS-CoV-2/human/USA/CA_CDC_5574/2020), and determined the relative abundance of each strain after replication and transmission by next generation sequencing. The results of this study confirm that adult white-tailed deer are highly susceptible to SARS-CoV-2 infection and shed virus in sufficient quantities through oral and nasal cavities to infect naïve contact sentinel deer. Furthermore, our results illustrate the *in vivo* competition of two lineages of SARS-CoV-2 through analysis of excreted virus and the virus presence in tissues collected *postmortem*. Importantly, this is the first study which provides evidence for vertical transmission of SARS-CoV-2 from doe to fetus.

## Materials and methods

### Cells and virus isolation/titrations

Vero E6 cells (ATCC; Manassas, VA) and Vero E6 cells stably expressing transmembrane serine protease 2 (Vero-E6/TMPRSS2)^7^ were obtained from Creative Biogene (Shirley, NY) via Kyeong-Ok Chang at KSU and used for virus propagation and titration. Cells were cultured in Dulbecco’s Modified Eagle’s Medium (DMEM, Corning, New York, N.Y, USA), supplemented with 5% fetal bovine serum (FBS, R&D Systems, Minneapolis, MN, USA) and antibiotics/antimycotics (ThermoFisher Scientific, Waltham, MA, USA), and maintained at 37 °C under a 5% CO_2_ atmosphere. The addition of the selection antibiotic, G418, to cell culture medium was used to maintain TMPRSS2 expression but was not used during virus cultivation or assays. The SARS-CoV-2/human/USA/WA1/2020 lineage A (referred to as lineage A WA1; BEI item #: NR-52281) and SARS-CoV-2/human/USA/CA_CDC_5574/2020 lineage B.1.1.7 (alpha VOC B.1.1.7; NR-54011) strains were acquired from BEI Resources (Manassas, VA, USA). A passage 2 plaque-purified stock of lineage A WA1 and a passage 1 of the alpha VOC B.1.1.7 stock were used for this study. Virus stocks were sequenced by next generation sequencing (NGS) using the Illumina MiSeq and the consensus sequences were found to be homologous to the original strains obtained from BEI (GISAID accession numbers: EPI_ISL_404895 (WA-CDC-WA1/2020) and EPI_ISL_751801 (CA_CDC_5574/2020).

To determine infectious virus titers of virus stocks and study samples, 10-fold serial dilutions were performed on Vero-E6/TMPRSS2 cells. The presence of cytopathic effects (CPE) after 96 hours incubation at 37 °C was used to calculate the 50% tissue culture infective dose (TCID_50_)/ml using the Spearman-Kaerber method^8^. Selected swab and tissue homogenate samples were tested for viable virus by culture on Vero E6/TMPRRS2 cells. Virus isolation was performed by culturing 400 μL of filtered (0.2 μm; MidSci, St. Louis, MO) sample on Vero E6 cells and monitoring for CPE for up to 5 days post inoculation. Virus isolation attempts were only performed on samples with ≥10^6^ RNA copy number per mL, as this was our approximate limit of detection (LOD) for viable virus using this method.

### Susceptibility of cervid cells to SARS-CoV-2

The SARS-CoV-2 USA-WA1/2020 strain was passaged 3 times in Vero-E6 cells to establish a stock virus for infection experiments. Primary white-tailed, mule deer and elk lung cells (provided by WCW) were infected at approximately 0.1 MOI. Infected cell supernatants were collected at 0, 2, 4, 6 or 8 days post infection (DPI) and stored at −80°C until further analysis. Cell lines were tested in at least two independent infection experiments. Cell supernatants were titrated on Vero E6 cells to determine TCID_50_/mL.

### Ethics statement

All animal studies and experiments were approved and performed under the Kansas State University (KSU) Institutional Biosafety Committee (IBC, Protocol #1460) and the Institutional Animal Care and Use Committee (IACUC, Protocol #4468) in compliance with the Animal Welfare Act. All animal and laboratory work were performed in biosafety level-3+ and −3Ag laboratories and facilities in the Biosecurity Research Institute at KSU in Manhattan, KS, USA.

### Virus challenge of animals

Six female white-tailed deer (WTD), approximately 2 years of age, were acquired from a Kansas deer farm (Muddy Creek Whitetails, KS) and acclimated for ten days in BSL-3Ag biocontainment with feed and water *ad libitum* prior to experimental procedures. On day of challenge, four principal infected deer were inoculated with a 1:1 titer ratio of lineage A WA1 and the alpha VOC B.1.1.7 strains (**Figure 1**). A 2 ml dose of 1×10^6^ TCID_50_ per animal was administered through intra-nasal (IN) and oral (PO) routes simultaneously. The remaining two non-infected deer were placed up-current of the room directional airflow from the principal infected deer, separated by an 8-foot tall, solid partition wall. At 1 day-post-challenge (DPC), the two naïve deer were co-mingled with the principal infected animals as contact sentinels for the duration of the study. Two principal infected deer were euthanized and *postmortem* examination performed at 4 DPC. *Postmortem* examination of the remaining two principal infected and two sentinels was performed at 18 DPC (**Table 1**). Five of the six deer were pregnant; the number of fetuses per deer are indicated in **Table 1**. Four naïve white-tailed deer from a previous study evaluating a baculovirus-expressed subunit vaccine for the protection from epizootic hemorrhagic disease (EHD), performed in 2017^9^, were used as controls (**Table 1 and Figure 1**).

**Figure 1.**
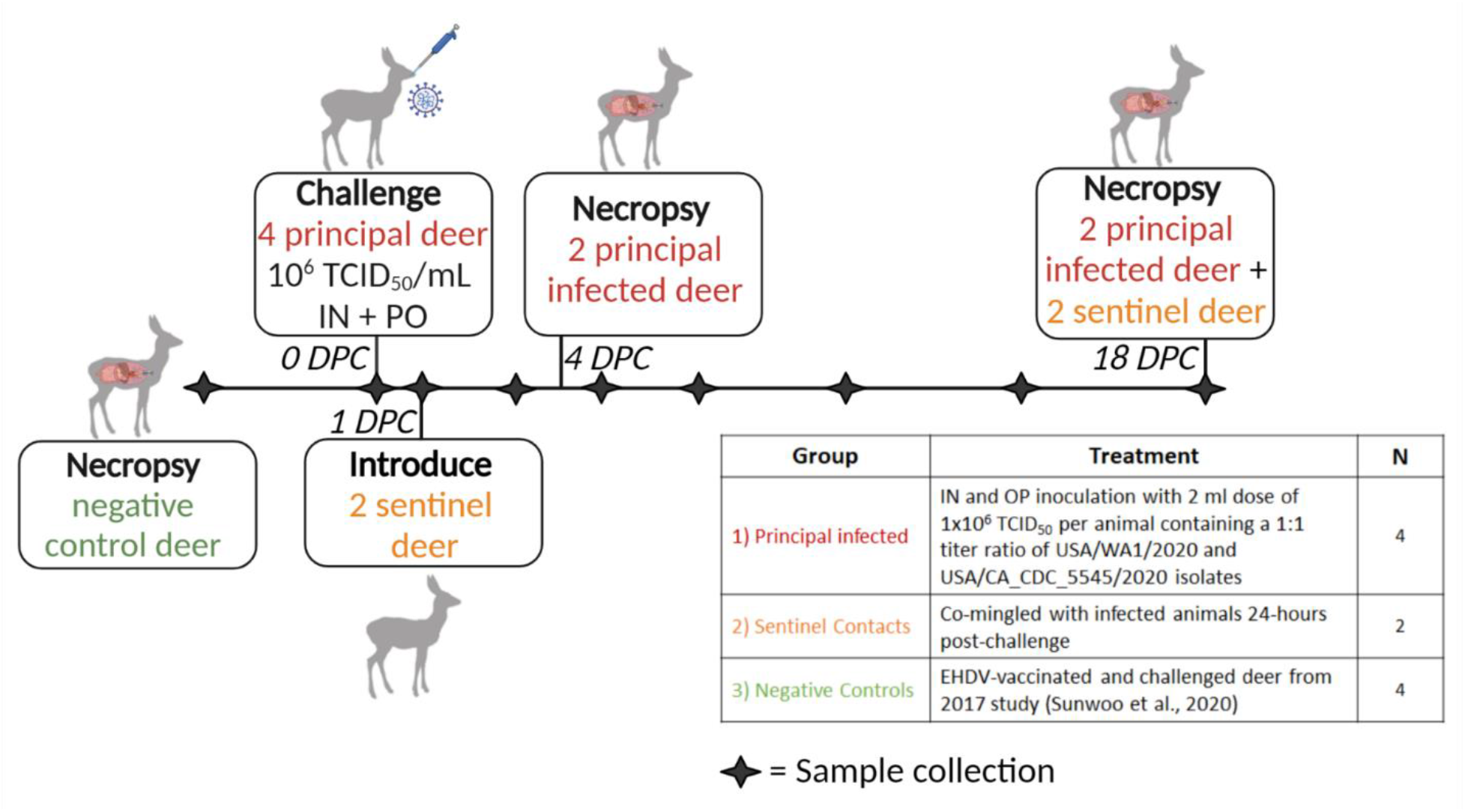
Experimental design. Ten female white-tailed deer were split into three groups as follows: *i)*four principal infected deer, *ii)* two sentinel contact deer, and *iii)* four non-inoculated control deer. Group 1 was inoculated simultaneously via intra-nasal and oral routes with a 2 ml dose of 1×10^6^ TCID_50_ per animal containing an approximate 1:1 titer ratio of the lineage A WA1 strain and an alpha VOC B.1.1.7 strain of SARS-CoV-2. Group 2 deer (n=2) were used as sentinel contact animals and were not challenged directly. These sentinel deer were placed up air current of the room’s directional airflow and separated from the principal infected group by an 8-foot tall, solid partition wall on the day of challenge, provided separate food and water, and re-introduced to principal infected (group 1) 24-hours post infection. Nasal, oral, and rectal swabs were collected on days 0, 1, 3, 5, 7, 10, 14, and 18 post-challenge. Whole blood and serum were collected on 0, 3, 7, 10, 14, and 18 DPC. Two principal infected deer (group 1) were euthanized for *postmortem* examination on 4 days-post-challenge (DPC) to evaluate the acute phase of infection. The four remaining deer, consisting of two sentinels and two principal infected, were maintained for the duration of the 18-day study to evaluate contact transmission and the convalescent stage of infection. The control deer were part of a separate study (36).

**Table 1.**
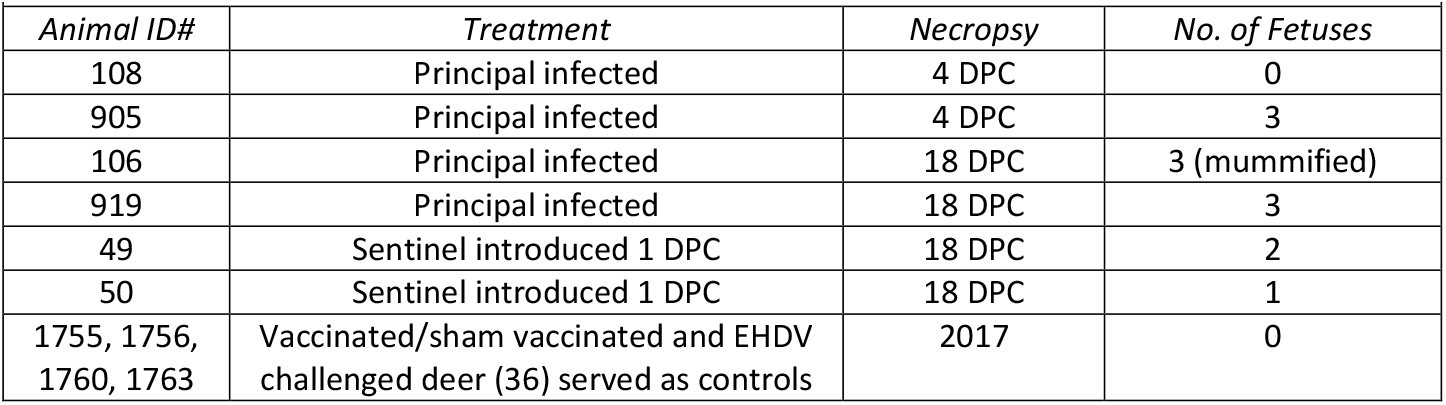
Animals and treatment assignments.

### Clinical evaluations and sample collection

Deer were observed daily for clinical signs. Clinical observations focused on activity level (response to human observer), neurological signs, respiratory rate, and presence of gastrointestinal distress. Rectal temperature, nasal, oropharyngeal, and rectal swabs were collected from sedated animals at 0, 1, 3, 5, 7, 10, 14 and 18 DPC. Swabs were placed in 2mL of viral transport medium (DMEM, Corning; combined with 1% antibiotic-antimycotic, ThermoFisher), vortexed, and aliquoted directly into cryovials and RNA stabilization/lysis Buffer RLT (Qiagen, Germantown, MD, USA). EDTA blood and serum were collected prior to challenge and on days 3, 7, 10, 14, and 18 DPC. Full *postmortem* examinations were performed, and gross changes recorded. A comprehensive set of tissues were collected in either 10% neutral-buffered formalin (Fisher Scientific, Waltham, MA, USA), or as fresh tissues directly stored at −80°C. Tissues were collected from the upper respiratory tract (URT) and lower respiratory tract (LRT), central nervous system (brain and cerebral spinal fluid [CSF]), gastrointestinal tract (GIT) as well as accessory organs. The lungs were removed *in toto* including the trachea, and the main bronchi were collected at the level of the bifurcation and at the entry point into the lung lobe. Lung lobes were evaluated based on gross pathology and collected and sampled separately. Bronchoalveolar lavage fluid (BALF), nasal wash and urine were also collected during *postmortem* examination. Fetal tissues including lung, liver, spleen, kidney as well as placenta were also collected. Fresh frozen tissue homogenates were prepared as described previously^10^. All clinical samples (swabs, nasal washes, BALF, CSF, urine) and tissue homogenates were stored at −80°C until further analysis.

### RNA extraction and reverse transcription quantitative PCR (RT-qPCR)

SARS-CoV-2 specific RNA was detected and quantified using a quantitative reverse transcription real time – PCR (RT-qPCR) assay specific for the N2 segment as previously described^11^. Briefly, nucleic acid extractions were performed by combining equal amounts of Lysis Buffer RLT (Qiagen, Germantown, MD, USA) with supernatant from clinical samples (swabs, nasal washes, BALF, CSF, urine), tissue homogenates in DMEM (20% W/V), EDTA blood or body fluids. Sample lysates were vortexed and 200 μL was used for extraction using a magnetic bead based extraction kit (GeneReach USA, Lexington, MA) and the Taco™ mini nucleic acid extraction system (GeneReach) as previously described^11^. Extraction positive controls (IDT, IA, USA; 2019-nCoV_N_Positive Control), diluted 1:100 in RLT lysis buffer, and negative controls were included throughout this process.

Quantification of SARS-CoV-2 RNA was accomplished using an RT-qPCR protocol established by the CDC for detection of SARS-CoV-2 nucleoprotein (N)-specific RNA (https://www.fda.gov/media/134922/download). Our lab has validated this protocol using the N2 SARS-CoV-2 primer and probe sets (CDC assays for RT-PCR SARS-CoV-2 coronavirus detection | IDT(idtdna.com)) in combination with the qScript XLT One-Step RT-qPCR Tough Mix (Quanta Biosciences, Beverly, MA, USA), as previously described^11^. Quantification of RNA copy number (CN) was based on a reference standard curve method using a 10-point standard curve of quantitated viral RNA (USA-WA1/2020; lineage A). This RT-qPCR assay was also validated to the detection of the USA/CA_CDC_5545/2020 alpha VOC B.1.1.7 strain. Each sample was run in duplicate wells and all 96-well plates contained duplicate wells of quantitated PCR positive control (IDT, IA, USA; 2019-nCoV_N_Positive Control, diluted 1:100) and four non-template control wells. A positive Ct cut-off of 38 cycles was used when both wells were positive. Samples with one of two wells positive at or under CT of 38 were considered suspect-positive. Data are presented as the mean and standard deviation of the calculated N gene CN per mL of liquid sample or per mg of 20% tissue homogenate.

### Next-Generation Sequencing

RNA extracted from cell culture supernatant (virus stocks), clinical swab/tissue homogenates, and clinical samples were sequenced by next generation sequencing (NGS) using an Illumina NextSeq platform (Illumina Inc.) to determine the genetic composition (% lineage) of viral RNA in each sample. SARS-CoV-2 viral RNA was amplified using the ARTIC-V3 RT-PCR protocol [Josh Quick 2020. nCoV-2019 sequencing protocol v2 (GunIt). Protocols.io https://gx.doi.org/10.17504/protocols.io.bdp7i5rn]. Library preparation of amplified SARS-CoV-2 DNA for sequencing was performed using a Nextera XT library prep kit (Illumina Inc.) following the manufacturer’s protocol. The libraries were sequenced on the Illumina NextSeq using 150 bp paired end reads with a mid-output kit. Reads were demultiplexed and parsed into individual sample files that were imported into CLC Workbench version 7.5 (Qiagen) for analysis. Reads were trimmed to remove ambiguous nucleotides at the 5’ terminus and filtered to remove short and low-quality reads. The consensus sequences of viral stocks used for challenge material preparation were found to be homologous to the original strains obtained from BEI (GISAID accession numbers: EPI_ISL_404895 (WA-CDC-WA1/2020) and EPI_ISL_751801 (CA_CDC_5574/2020). To determine an accurate relative percentage of each SARS-CoV-2 lineage in each sample, BLAST databases were first generated from individual trimmed and filtered sample reads. Subsequently, two 40-nucleotide long sequences were generated for each strain at locations that include the following gene mutations: Spike (S) A570D, S D614G, S H1118H, S ΔH69V70, S N501Y, S P681H, S 982A, S T716I, S ΔY145, Membrane (M) V70, Nucleoprotein (N) D3L, N R203KG204R, N 235F, and non-structural NS3 T223I, NS8 Q27stop, NS8 R52I, NS8 Y73C, NSP3 A890D, NSP3 I1412T, NSP3 T183I, NSP6 ΔS106G107F108, NSP12 P323L, NSP13 A454V and NSP13 K460R. A word size of 40 was used for the BLAST analysis in order to exclude reads where the target mutation fell at the end of a read or reads that partially covered the target sequence. The two sequences for each of the locations listed above, corresponding to either lineage A (USA-WA1/2020) or alpha VOC B.1.1.7 (USA/CA_CDC_5574/2020), were subjected to BLAST mapping analysis against individual sample read databases to determine the relative amount of each strain in the samples. The relative amount of each strain present was determined by calculating the percent of reads that hit each of the twenty-three target mutations. The percentages for all mutations were averaged to determine the relative amount of each strain in the sample. Samples with incomplete or low coverage across the genome were excluded from analysis. Average depth of read coverage ranged from 554 up to 48315 with 9312 as the median coverage per sample (**Supplementary Figure 1**).

### Virus neutralizing antibodies

Virus neutralizing antibodies in sera were determined using microneutralization assay as previously described^11^. Briefly, heat inactivated (56°C/ 30 min) serum samples were subjected to 2-fold serial dilutions starting at 1:20 and tested in duplicate. Then, 100 TCID_50_ of SARS-CoV-2 virus in 100 μL DMEM culture media was added 1:1 to 100 μL of the sera dilutions and incubated for 1 hour at 37°C. The mixture was subsequently cultured on Vero-E6/TMPRSS2 cells in 96-well plates. The neutralizing antibody titer was recorded as the highest serum dilution at which at least 50% of wells showed virus neutralization based on the absence of CPE observed under a microscope at 72 h post infection.

### Detection of antibodies by indirect ELISA

Indirect ELISAs were used to detect SARS-CoV-2 antibodies in sera with nucleocapsid (N) and the receptor-binding domain (RBD) recombinant viral proteins, both produced in-house^11^. Briefly, wells were coated with 100 ng of the respective protein in 100 μL per well coating buffer (Carbonate–bicarbonate buffer, catalogue number C3041, Sigma-Aldrich, St. Louis, MO, USA). Following an overnight incubation at 4 °C, plates were washed three times with phosphate buffered saline (PBS-Tween 20 [pH=7.4]; catalogue number 524653, Millipore Sigma), blocked with 200 μL per well casein blocking buffer (Sigma-Aldrich, catalogue number B6429) and incubated for 1 hour at room temperature (RT). Plates were subsequently washed three times with PBS-Tween-20 (PBS-T). Serum samples were diluted 1:400 in casein blocking buffer, then 100 μL per well was added to ELISA plates and incubated for 1 hour at RT. Following three washes with PBS-T, 100 μL of HRP-labelled Rabbit Anti-Deer IgG (H+L) secondary antibody (95058-328, VWR, Batavia, IL, USA) diluted 1:1000 (100ng/mL) was added to each well and incubated for 1 hour at RT. Plates were then washed five times with PBS-T and 100 μL of TMB ELISA Substrate Solution (Abcam, catalogue number ab171525, Cambridge, MA, USA) was added to all wells of the plate and incubated for 5 minutes before the reaction was stopped. The OD of the ELISA plates were read at 450 nm on an ELx808 BioTek plate reader (BioTek, Winooski, VT, USA). The cut-off for a sample being called positive was determined as follows: Average OD of negative serum + 3X standard deviation. Everything above this cut-off was considered positive. Indirect ELISA was used to detect Bovine Coronavirus (BCoV) antibodies in sera with Spike (S) recombinant viral protein (LSBio, LS-G64076-20, Seattle, WA, USA) using the methods described above.

### Histopathology

Tissue samples from the respiratory tract (nasal cavity [rostral, middle and deep turbinates following decalcification with Immunocal™ Decalcifier (StatLab, McKinney, TX, for 4-7 days at room temperature), trachea, and lungs as well as various other extrapulmonary tissues (liver, spleen, kidneys, heart, pancreas, gastrointestinal tract [stomach, small intestine including Peyer’s patches and colon], cerebrum [including olfactory bulb], tonsils and numerous lymph nodes were routinely processed and embedded in paraffin. Four-micron tissue sections were stained with hematoxylin and eosin following standard procedures. Two independent veterinary pathologists (blinded to the treatment groups) examined the slides and morphological descriptions were provided.

### SARS-CoV-2-specific immunohistochemistry (IHC)

IHC was performed as previously described^11^ on four-micron sections of formalin-fixed paraffin-embedded tissue mounted on positively charged Superfrost® Plus slides and subjected to IHC using a SARS-CoV-2-specific anti-nucleocapsid rabbit polyclonal antibody (3A, developed by our laboratory) with the method previously described^12^. Lung sections from a SARS-CoV-2-infected hamster were used as positive assay controls.

## Results

### Susceptibility of cervid primary lung cells to SARS-CoV-2

Primary lung cells isolated from white-tailed deer, mule deer and elk were tested for susceptibility to SARS-CoV-2 and viral growth kinetics. SARS-CoV-2 lineage A WA1 strain was found to replicate in both, white-tailed deer and mule deer lung cells but not in elk lung cells (**Figure 2A**), and replication resulted in cell death (**Figure 2B**). Similar kinetics were observed at 2 and 4 DPI for both the white-tailed deer and mule deer cells. Virus titers declined by 6 DPC in white-tailed deer lung cells, while titers in mule deer lung cultures remained elevated even at 8 DPC. Elk lung cell input virus rapidly declined by day 2 and virus was not detectable by 8 DPC. These results suggest that besides white-tailed deer, mule deer species may be susceptible to SARS-CoV-2 and could serve as a potential reservoir species.

**Figure 2.**
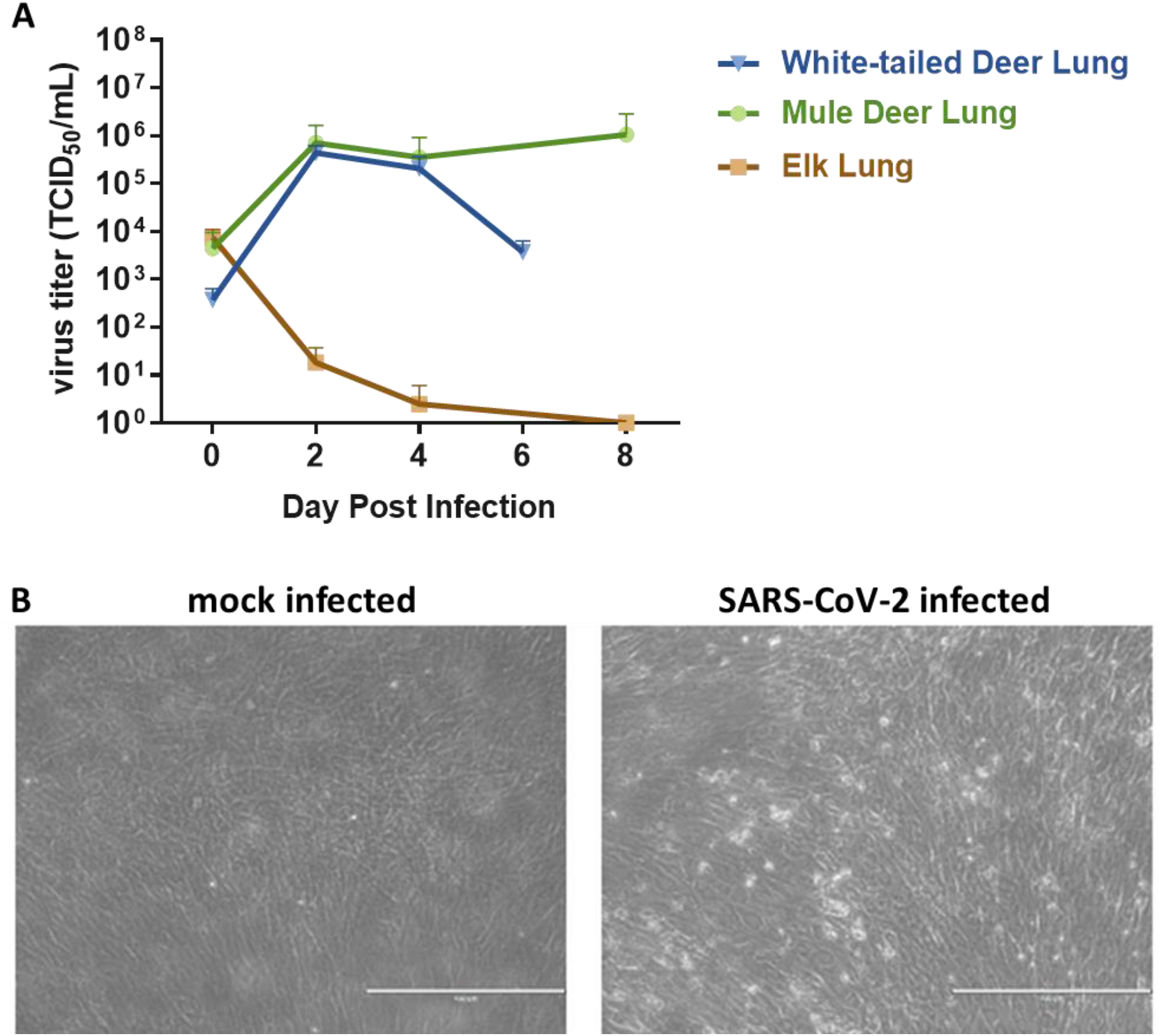
SARS-CoV-2 replication in various cervid lung cells. (**A**) Primary lung cells were infected with the SARS-CoV-2 USA-WA1/2020 at 0.1 MOI and cell supernatants collected at 0, 2, 4, 6 or 8 days post infection (DPI). Cell supernatants were titrated on Vero E6 cells to determine virus titers. Mean titers of at least two independent infection experiments per cell line are shown. (**B**) Cytopathic effect observed at 6 DPI with SARS-CoV-2 but not in mock infected white-tailed deer primary lung cells at same time point DPI.

### SARS-CoV-2-infected adult white-tailed deer remain subclinical

Four deer were inoculated with an approximate 1:1 titer ratio of the lineage A WA1 and alpha VOC B.1.1.7 strains (**Figure 1**) with 1×10^6^ TCID_50_ per animal administered through IN and PO routes simultaneously. Clinical signs were recorded daily, including respiratory rate, posture, and activity levels. Rectal temperatures were recorded on days of sample collection. No major clinical signs were observed throughout the course of this study. Ocular discharge was noted in the principal infected animal #106 from 5 to 10 DPC and nasal discharge was noted on 7 DPC in principal infected animal #919. Rectal temperatures of principal and sentinel deer remained within normal range (99-105°F), although there was a slight elevation in body temperature of principal infected deer from 0 to 3 DPC, and at 1 to 3 DPC in the sentinels (**Supplementary Figure 2**). Soft stool was observed in one deer (#905) for the duration of the study; this was not considered as a result from SARS-CoV-2 infection as this was also observed prior to challenge.

### SARS-CoV-2 RNA/virus shedding

SARS-CoV-2 RNA shedding was determined by evaluating nasal, oral and rectal swabs of principal infected (**Figure 3A**) and sentinel deer (**Figure 3B**) for the presence of SARS-CoV-2 specific RNA by RT-qPCR. Viral RNA was detected in nasal swabs from all principal infected animals (n=4) from 1 to 10 DPC (**Figure 3A**), and the sentinel contact deer (n=2) starting at 3 DPC (2 days post comingling) up to 10 DPC (**Figure 3B**). All deer had SARS-CoV-2 RNA detected in oral swabs at some time point post infection/contact, although less frequently and at lower levels compared to nasal swabs. Viral RNA from oral swabs was detected from 1 to 7 DPC in at least one of the principal infected deer. Oral swabs from at least one sentinel deer were detectable at 3 and 5 DPC (2 to 4 days post comingling), and both sentinels were positive at 7 DPC. Rectal swabs had low levels of viral RNA detected at 5 and 7 DPC in two principal infected animals, and in only one sentinel deer at 5 DPC. Infectious virus could be isolated from nasal swabs of at least one of the principal infected deer at 1, 3 and 4 DPC, and from one sentinel at 7 DPC (**Table 2**). Infectious virus was isolated from oral swabs from one principal infected animal at 3 DPC and from one sentinel at 5 DPC (**Table 2**). In addition, virus was also isolated from a single rectal swab at 5 DPC from a principal infected deer.

**Figure 3.**
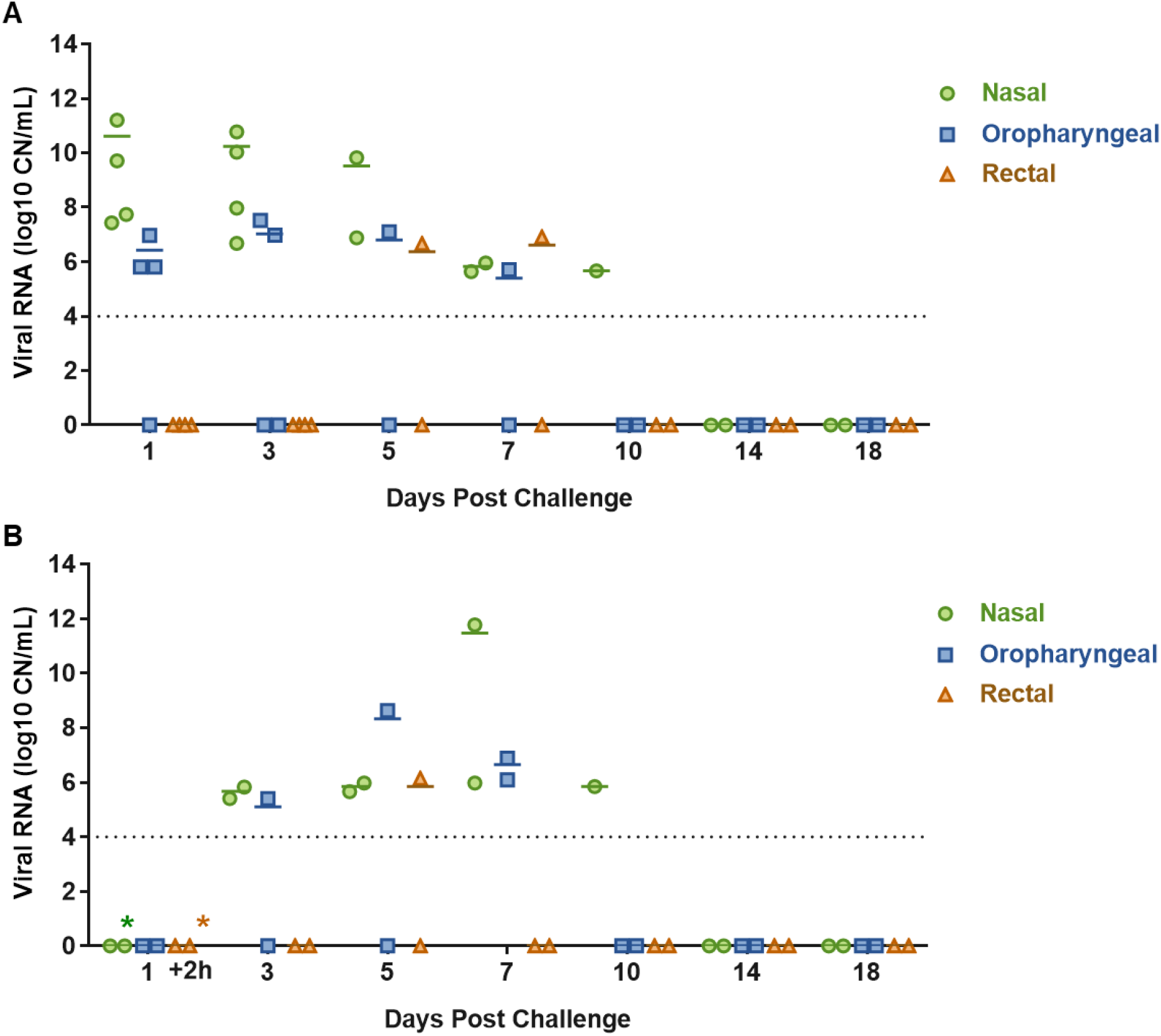
Viral shedding of SARS-CoV-2-infected white-tailed deer. RT-qPCR was performed on nasal, oropharyngeal, and rectal swabs collected from principal infected **(A)** and sentinel deer **(B)** on the indicated days post challenge (DPC). Mean (n=2) viral RNA copy number (CN) per mL of the SARS-CoV-2 nucleocapsid gene is reported. The limit of detection for this assay is indicated by the dotted lines. Asterisks (*) indicate samples with 1 out of 2 RT-qPCR reactions below the limit of detection.

**Table 2.** Virus isolation from swabs and tissues.

| **Table 2. Virus isolation from swabs and tissues** |  |  |  |
| --- | --- | --- | --- |
| *Sample Type* | *DPC* | *Animal Number* | *Virus titer (TCID<sub>50</sub>/mL)* |
| Nasal Swab | 1 | #106, 905, 919 | NEG |
| Nasal Swab | 1 | #108 | 1.00E+02 |
| Nasal Swab | 3 | #106 | 9.98E+02 |
| Nasal Swab | 3 | #108, 905, 919 | NEG |
| Nasal Swab | 4 | #108 | 9.98E+02 |
| Nasal Swab | 4 | #905 | NEG |
| Nasal Swab | 5 | #106, 919 | NEG |
| Nasal Swab | 7 | #49 | 3.15E+02 |
| Oral Swab | 1 | #905 | NEG |
| Oral Swab | 3 | #106 | 5.00E+00 |
| Oral Swab | 3 | #108 | NEG |
| Oral Swab | 4 | #108, 905 | NEG |
| Oral Swab | 5 | #50 | 5.00E+00 |
| Oral Swab | 5 | #106 | NEG |
| Oral Swab | 7 | #49, 50 | NEG |
| Rectal Swab | 5 | #49, 919 | NEG |
| Rectal Swab | 5 | #106 | 3.15+e02 |
| Rectal Swab | 7 | #919 | NEG |
| Nasopharynx | 4 | #108 | NEG |
| Conche | 4 | #108 | NEG |
| Ethnoturbinates | 4 | #108 | NEG |
| Trachea | 4 | #108 | 3.15E+02 |
| Bronchi | 4 | #108 | 3.15E+02 |
| Lung | 4 | #108 | NEG |
| Tonsil | 4 | #108, 905 | NEG |
| Lymph node | 4 | #108, 905 | NEG |
| CSF | 4 | #108 | NEG |
| BALF | 4 | #108, 905 | 3.15E+03, NEG |
| Nasal Wash | 4 | #108, 905 | 3.15E+02, NEG |
| Nasopharynx | 18 | #50, 919 | NEG |
| Conche | 18 | #50 | NEG |
| Ethnoturbinates | 18 | #50 | NEG |
| Tonsil | 18 | #50 | NEG |
| Lymph node | 18 | #50 | NEG |
| Bone Marrow | 18 | #50 | NEG |
| *Only samples with >10<sup>6</sup> RNA copy number were subjected to virus isolation.* |  |  |  |

### Evaluation of deer during the acute stage of infection (4 DPC)

To evaluate the acute stage of infection, two of the principal infected deer (#108 and #905) were humanely euthanized and examined at 4 DPC. SARS-CoV-2 RNA was detected throughout the upper and lower respiratory tract tissues of both deer necropsied at 4 DPC (**Figure 4A**). The highest levels of SARS-CoV-2 RNA were detected in the nasal cavity, trachea, bronchi, and all lung lobe sections. There was a distinct difference in viral RNA load observed in the respiratory tract tissues between the two deer at 4 DPC; one deer (#905) revealed consistently lower quantities of SARS-CoV-2 RNA throughout upper and lower respiratory tissues than the other deer (#108). Nasal washes and BALF from both animals were also positive for SARS-CoV-2 RNA at 4 DPC. Infectious virus was isolated from the nasal wash, BALF, trachea and bronchi of deer #108, but not from any tissues tested for deer #905 (**Table 2**).

**Figure 4.**
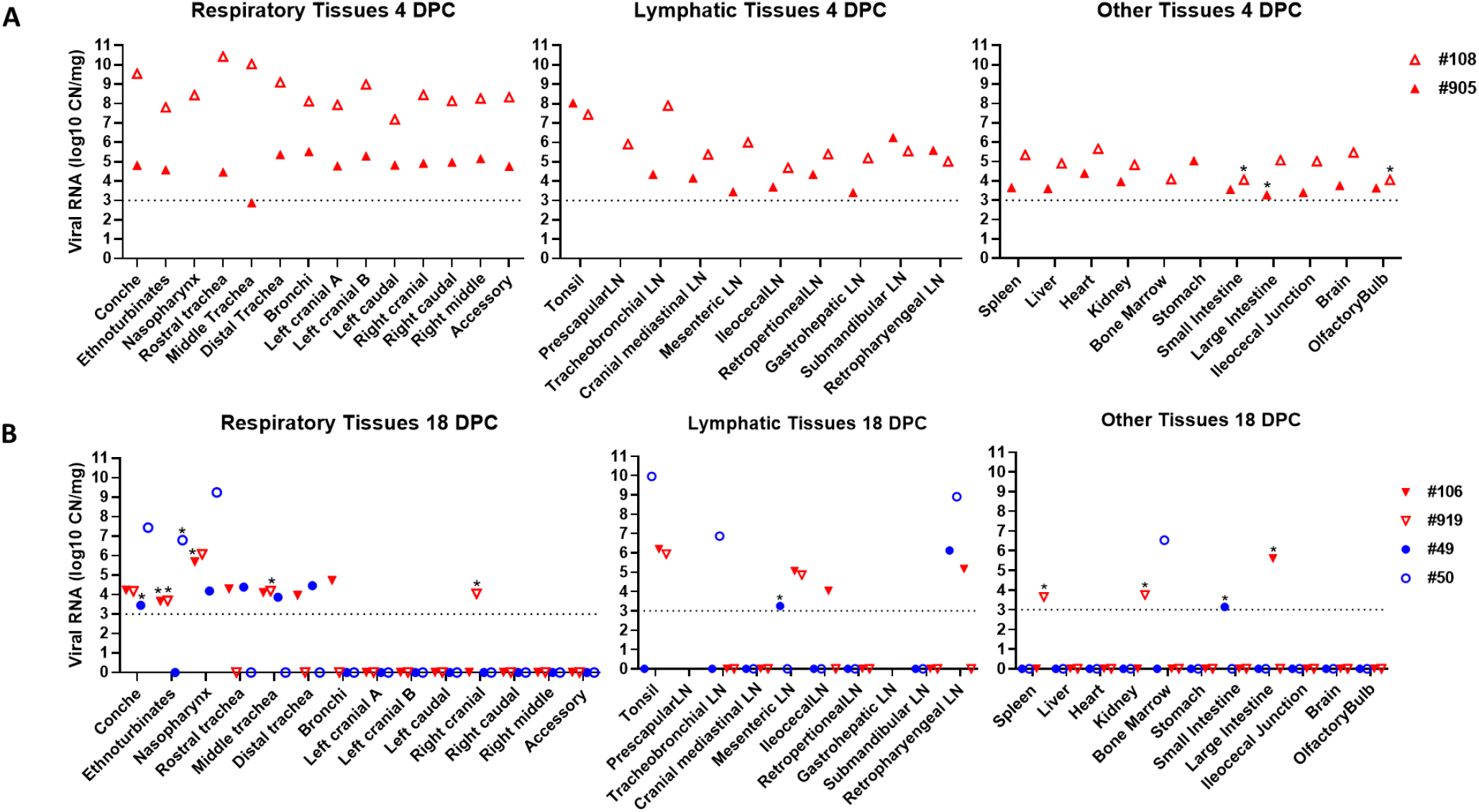
SARS-CoV-2 RNA Detected in Tissues. RT-qPCR was used to detect the presence of SARS-CoV-2 specific RNA in various tissues of deer euthanized at 4 (**A**) and 18 days (**B**) post challenge (DPC). Mean (n=2) viral RNA copy number (CN) per mg based on the nucleocapsid gene are plotted for individual animals. Colored symbols correspond to deer ID numbers: red triangles for principal infected deer and blue circles for sentinels. Asterisks (*) indicate samples with 1 out of 2 RT-qPCR reactions below the limit of detection, which is indicated by the dotted lines.

Histological evaluations and IHC for the presence of SARS-CoV-2 antigen in the upper and lower respiratory tract tissues, tonsils and select lymph nodes collected at 4 DPC were performed. In the respiratory tract, histological changes were more pronounced in deer #108 compared to #905, and a mild to moderate multifocal lymphohistiocytic and neutrophilic rhinitis, erosive to suppurative tracheitis and erosive bronchitis with mixed peribronchiolitis were noted (**Figure 5**). Specifically, there was marked attenuation of the respiratory epithelium lining the trachea and a main bronchus with loss of cilia, individual cell degeneration and necrosis, neutrophil transmigration, and accumulation of cellular debris in the lumen. The subjacent edematous lamina propria was infiltrated by neutrophils, lymphocytes and histiocytes; and respiratory epithelial cells frequently contained intracytoplasmic viral antigen, also abundant in the superficial exudate. In the remaining pulmonary parenchyma, bronchioles and blood vessels were delineated by perivascular and peribronchiolar lymphocytes, histiocytes and few neutrophils, but viral antigen was generally not detected. Rarely, sloughed and necrotic epithelial cells and few degenerate leukocytes lodged at the termini of respiratory bronchioles and containing intracytoplasmic viral antigen were observed (**Figure 5**). In deer #905, the inflammatory component was mild and predominantly lymphocytic with no viral antigen detected in the respiratory tract. Along the rostral turbinates of both animals (#905 and 108), the lamina propria was infiltrated by lymphocytes, histiocytes and fewer neutrophils that transmigrated through the lining nasal epithelium and encircled subjacent nasal glands. In contrast, the deep turbinates including the olfactory neuroepithelium, olfactory nerve fascicles and olfactory bulb were unremarkable. No viral antigen was detected within the segments of the nasal passages that were evaluated (**Figure 5**). A mild erosive tonsilitis with a small focus of intraepithelial SARS-CoV-2 antigen was also noted in deer #108 at 4 DPC (**Supplementary Figure 3**). No other significant histologic alterations were found in other tissues examined from the adult deer. Fetal lungs occasionally contained intrabronchial/intrabronchiolar squames; however, no viral antigen was detected here or within the placenta.

**Figure 5.**
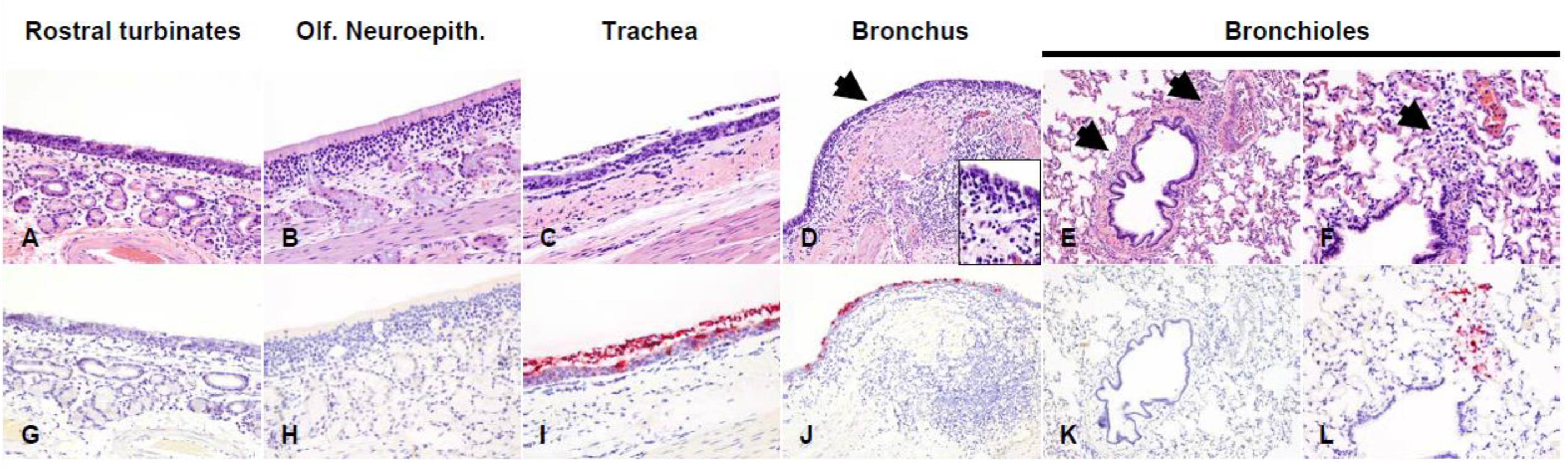
Histological lesions and SARS-CoV-2 antigen distribution in the upper and lower respiratory tract of principal infected white-tailed deer at 4 DPC. Rostral turbinates (**A** and **G**), olfactory neuroepithelium (**B** and **H**), trachea (**C** and **I**), bronchus (D and J) and bronchioles (E, F, K and L). At 4 DPC, in the rostral turbinates, neutrophilic rhinitis with epithelial transmigration and mixed lymphocytic and histiocytic infiltration of the subjacent lamina propria and around nasal glands were observed (**A**) but no viral antigen was detected (**G**). The olfactory neuroepithelium was histologically unremarkable (**B**) with no viral antigen detected (**H**). In the trachea, there was marked attenuation of the respiratory epithelium with loss of cilia, individual cell degeneration and necrosis, neutrophil transmigration, and accumulation of cellular debris in the lumen (**C**). Frequently, respiratory epithelial cells of the trachea contained intracytoplasmic viral antigen, which was also abundant in the superficial exudate (**I**). The bronchial mucosa was characterized by segmental attenuation of the lining respiratory epithelium with loss of cilia, degeneration/necrosis of individual epithelial cells and neutrophil and lymphocyte transmigration, and a mixed lymphocytic and histiocytic infiltrate in the edematous lamina propria (**D**, arrows and inset). The bronchial epithelium lining affected segments frequently contained viral antigen (**J**). In the pulmonary parenchyma, bronchioles and blood vessels were delimited by perivascular and peribronchiolar lymphocytes, histiocytes and few neutrophils (**E**, arrows). Viral antigen was generally not detected (**K**). In bronchioles, rarely sloughed and necrotic epithelial cells and few degenerate leukocytes lodged at the termini of respiratory bronchioles (**F**, arrow) contained intracytoplasmic viral antigen (**L**). H&E and Fast Red, 100X total magnification.

### Evaluation of deer during the convalescent stage of infection (18 DPC)

*Postmortem* examination of the remaining four deer, consisting of two principal infected animals and two contact sentinels, was performed on 18 DPC. SARS-CoV-2 RNA was detected in the upper respiratory tract tissues (nasal cavity and trachea) of the principal infected and sentinel deer at 18 DPC (**Figure 4B**). One of the principal infected animals had viral RNA in the bronchi. The lung lobes were negative for viral RNA in all deer except for the right cranial lung lobe from one of the principal infected deer (#919) that was considered as suspect. No viral RNA was detected from nasal wash or BALF collections from any of the deer at 18 DPC. Infectious virus was not detected in different tissues from one sentinel animal (#50) and in the nasopharynx of one principal infected animal (#919; **Table 2**).

Histologic examination at 18 DPC of principal infected deer #106 and #919 showed minimal changes in the lungs, with scattered peribronchiolar/perivascular cuffs of lymphocytes and plasma cells (**Figure 6**). Principal infected deer #919 showed evidence of chronic suppurative bronchopneumonia restricted to one lung lobe, but the overall histological changes suggest a possible underlying bacterial component (data not shown). Minimal mononuclear inflammation was noted in the trachea. Similarly, deer #106 had minimal changes in the trachea with rare mononuclear inflammatory aggregates within the lamina propria and extending into the lining respiratory epithelium (**Figure 6**). No viral antigen was detected by IHC in any of the respiratory tissues of principal infected deer at 18 DPC. The sentinel contact animals (#49 and #50) showed predominantly histological alterations along the trachea, with mild to moderate lymphoplasmacytic and erosive tracheitis characterized by multifocal segments of epithelial attenuation, necrosis and loss with occasional areas of epithelial hyperplasia, intense lymphoplasmacytic and neutrophilic inflammation along the superficial lamina propria, and luminal necrotic cell debris (**Figure 6**). While the changes in the lungs were mostly characterized by minimal peribronchiolar lymphocytic cuffing, deer #49 had a localized area of subacute to chronic suppurative bronchopneumonia with evident intralesional bacteria, likely representing a secondary opportunistic infection. Moderate lymphoplasmacytic pharyngitis with occasional attenuation and loss of surface epithelium and lymphoid follicle formation, and marked tonsillar lymphoid hyperplasia were also noted (data not shown). No other significant histologic alterations were identified in principal or sentinel animals, including their fetuses. Viral antigen was neither detected in the respiratory tract tissues (**Figure 6**) nor in placenta or fetal tissues derived from sentinel animals.

**Figure 6.**
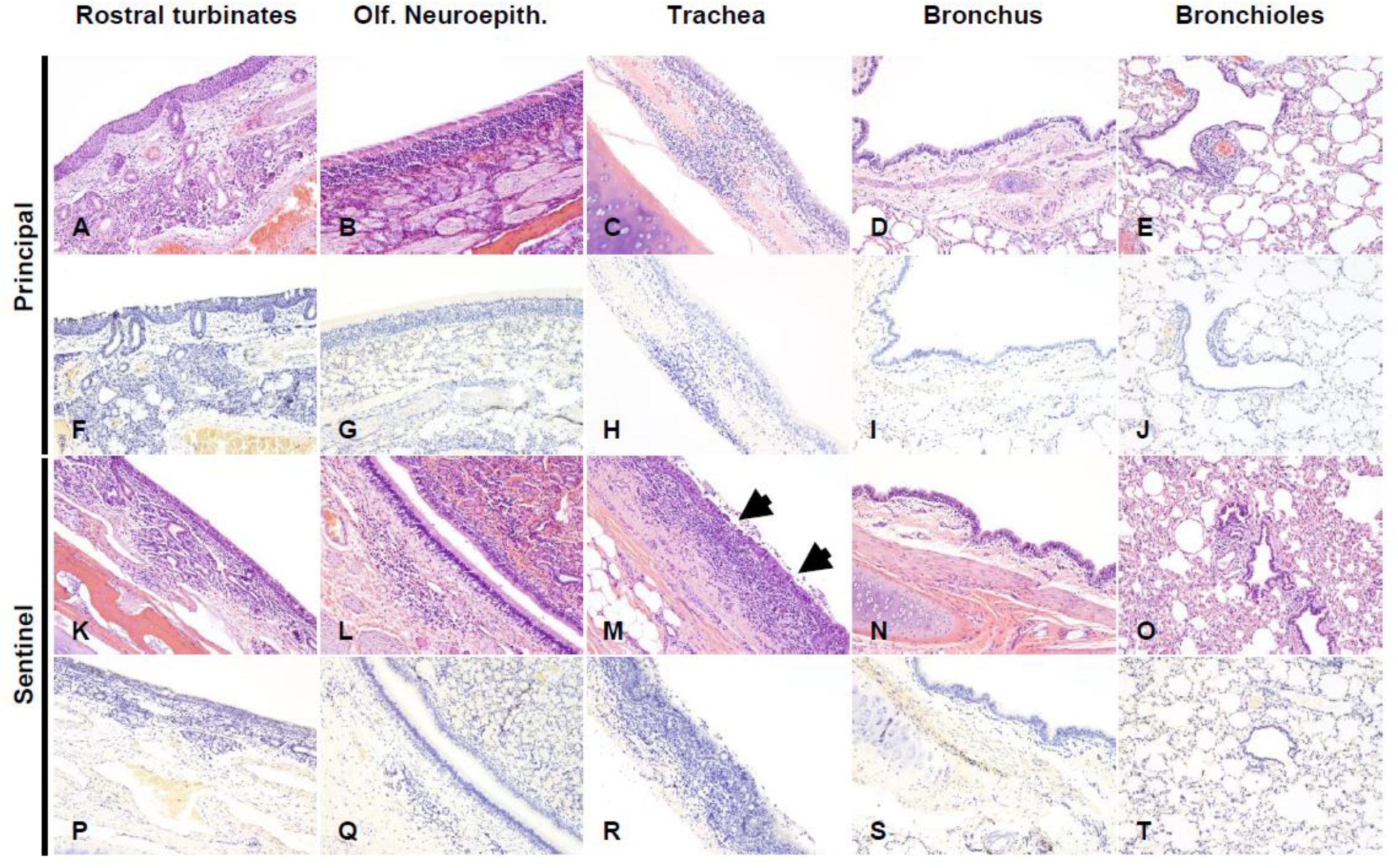
Histological lesions and SARS-CoV-2 antigen distribution in the upper and lower respiratory tract of principal infected and sentinel white-tailed deer at 18 DPC. Rostral turbinates (**A, F, K, P**), olfactory neuroepithelium (**B, G, L, Q**), trachea (**C, H, M, R**), bronchus (**D, I, N, S**) and bronchioles (**E, J, O, T**). In principal infected deer, few lymphocytes were noted in the lamina propria of the rostral turbinates (**A**). The olfactory neuroepithelium was histologically unremarkable (**B**), and the tracheal and bronchial lamina propria were occasionally infiltrated by either dispersed or aggregates of mononuclear cells (**C, D**), which also encircled few bronchioles and pulmonary vessels (**E**). In sentinel deer, the lamina propria of rostral turbinates and sporadically subjacent to the olfactory neuroepithelium were infiltrated by mild numbers of lymphocytes and plasma cells (**K, L**). There was segmental erosive tracheitis with epithelial attenuation and necrosis, and intense lymphoplasmacytic and neutrophilic inflammation (**M**, arrows). Few mononuclear cells were noted delimiting bronchi, bronchioles and pulmonary vessels (**N, O**). No viral antigen was detected in respiratory tract tissues of principal infected (**F-J**) and sentinel (**P-T**) deer at 18 DPC. H&E and Fast Red, 100X total magnification.

### Presence of SARS-CoV-2 RNA in non-respiratory organs and tissues during the acute and convalescent stage of infection

At 4 DPC, tissues collected from the two principal infected deer demonstrate a systemic presence of viral RNA (**Figure 4A**). Both deer had especially high levels of viral RNA detected in the tonsil. Lymph nodes from both animals, which included the cranial mediastinal, retropharyngeal, submandibular, tracheobronchial, prescapular, mesenteric, ileocecal, retroperitoneal, and gastrohepatic, were all positive for SARS-CoV-2 RNA at 4 DPC. Other tissues consisting of spleen, liver, heart, kidney, bone marrow, stomach, ileocecal junction and brain were also all positive for viral RNA in both animals at 4 DPC. Small and large intestine and olfactory bulb tissues were collected from the two principal infected deer at 4 DPC, and these tissues were RT-qPCR positive in one of the two animals whereas the other one was considered suspect. CSF obtained from deer #108 was also positive for SARS-CoV-2 RNA at 4 DPC.

At 18 DPC, SARS-CoV-2 RNA was detected at relatively high levels in the tonsil of both principal infected and one sentinel animal (**Figure 4B**). Several lymph nodes were also positive for viral RNA at 18 DPC, although fewer compared to 4 DPC, and included the mesenteric, ileocecal and retropharyngeal lymph nodes from principal infected deer, and the tracheobronchial, retropharyngeal and a suspect positive mesenteric lymph node from the sentinels. At 18 DPC, SARS-CoV-2 RNA was not detected in the liver, heart, stomach, brain or olfactory bulb. The spleen, kidney and large intestine of at least one principal infected deer, and the small intestine of one sentinel were suspect positive for viral RNA at 18 DPC. No viral RNA was detected in the CSF or urine from any deer at 18 DPC. In addition, no infectious virus was isolated in samples collected from one principal infected (#919) and one sentinel (#50) at 18 DPC (**Table 2**).

### SARS-CoV-2-specific markers present in deer fetuses

Fetal tissues were collected from 5 pregnant deer consisting of 12 fetuses in total – one animal (#108) was not pregnant (**Table 1**). Full tissue sample sets, consisting of spleen, liver, kidney, lungs, and placenta were collected when possible (from 6 out of the 12 fetuses). Two of the three fetuses from deer #905 collected at 4 DPC, had detectable amounts of SARS-CoV-2 RNA in at least one of the tissues collected: one fetus was RNA positive in the lungs, liver, spleen, and placenta while the other fetus had viral RNA detected only in the spleen (**Table 3**). At 18 DPC, partial tissue sample sets were collected from the nine fetuses present in the four remaining pregnant deer (**Table 1**); they were all negative for SARS-CoV-2 RNA (**Table 3**). The three fetuses from deer #106 were found mummified, two viable fetuses each were obtained from deer #919 and #49, whereas the remaining two fetuses analyzed at 18 DPC appeared to be non-viable. This indicates that more than 50% (5/9) of the fetuses collected on 18 DPC were non-viable. Only the uterus of deer #108 which was not pregnant was positive for SARS-CoV-2 RNA at 4 DPC. The two RNA-positive fetuses from doe #905 were negative for the presence of SARS-CoV-2 antigen by IHC.

**Table 3.**
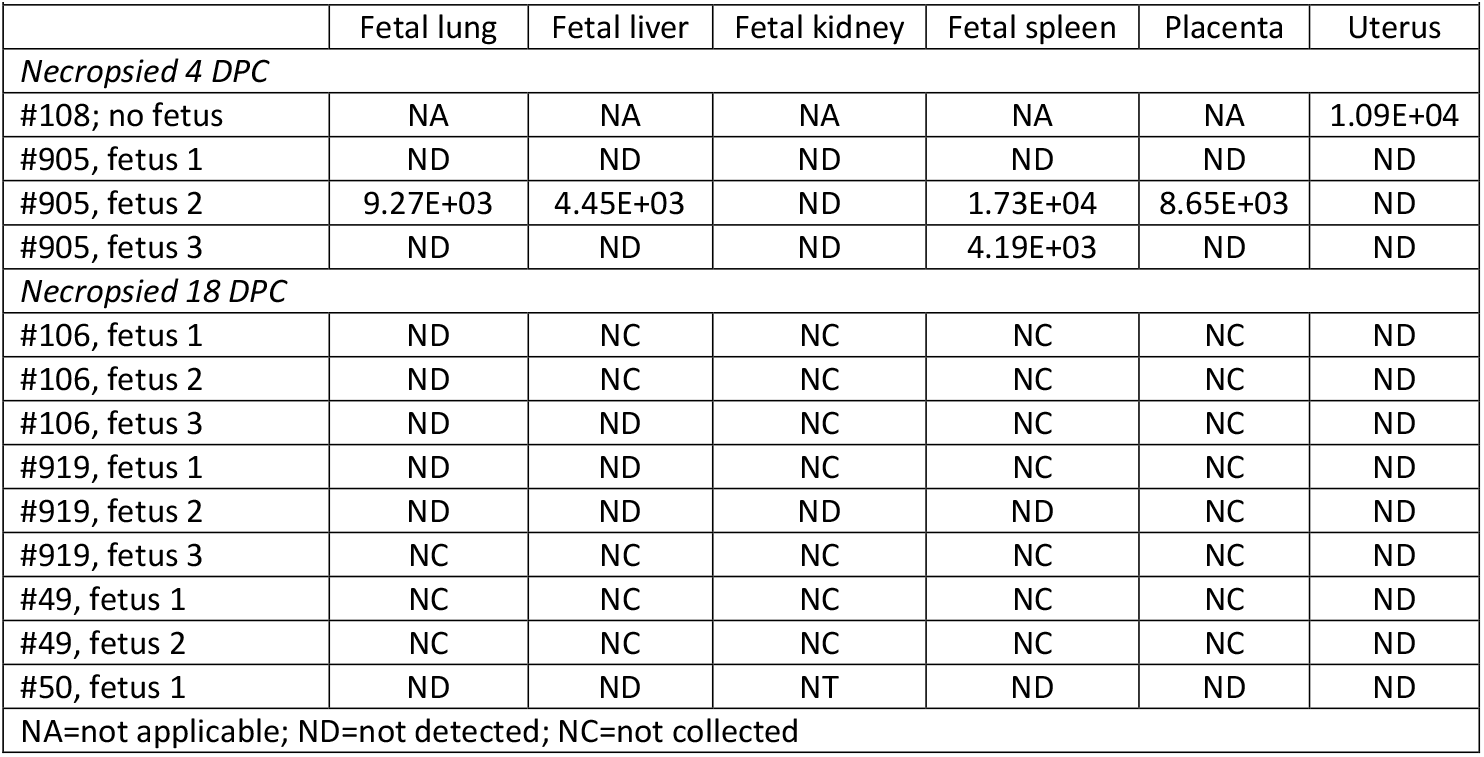
Presence of SARS-CoV-2 RNA (CN/mg) in fetuses and associated tissues.

### Serology

SARS-CoV-2-specific and neutralizing serum antibodies were evaluated for control, principal infected and sentinel deer (**Figure 7**). Some of the control sera which includes pre-challenge sera (−3 DPC) were weak positive in the ELISA for the SARS-CoV-2 nucleocapsid protein N. The principal infected animals and the sentinels remained weak positive over the 18-day course of the study (**Figure 7A**); only one principal infected deer developed substantial N-specific antibodies at 7, 14 and 18 DPC. Similarly, some of the control sera were weak positive in the ELISA for the SARS-CoV-2 RBD antigen. The two principal infected animals remaining after 4 DPC were clearly positive at 7 DPC with subsequent lower OD levels observed at 14 and 18 DPC, whereas the sentinel deer remained weak positive or even negative for RBD-specific antibodies over the 18-day course of the study (**Figure 7B**). The presence of virus neutralizing antibodies in the sera was tested in a classical virus neutralization assay. Some of the control sera had borderline levels of reactivity in the virus neutralization assay. However, at 7 DPC, significant levels of neutralizing antibody titers were observed in sera of principal infected deer, and similarly in the sera of sentinel deer at 10 DPC (**Figure 7C**). At 14 and 18 DPC, both the principal infected and sentinel deer had similar titers (approximately 1:1280) of neutralizing antibodies (**Figure 7C**). Finally, sera from the principal infected and sentinel deer were tested for the presence of bovine coronavirus S-specific antibodies by an indirect ELISA. Although the OD value was slightly higher in some of the principal infected animals, all were at near negative bovine control sera levels (**Figure 7D**).

**Figure 7.**
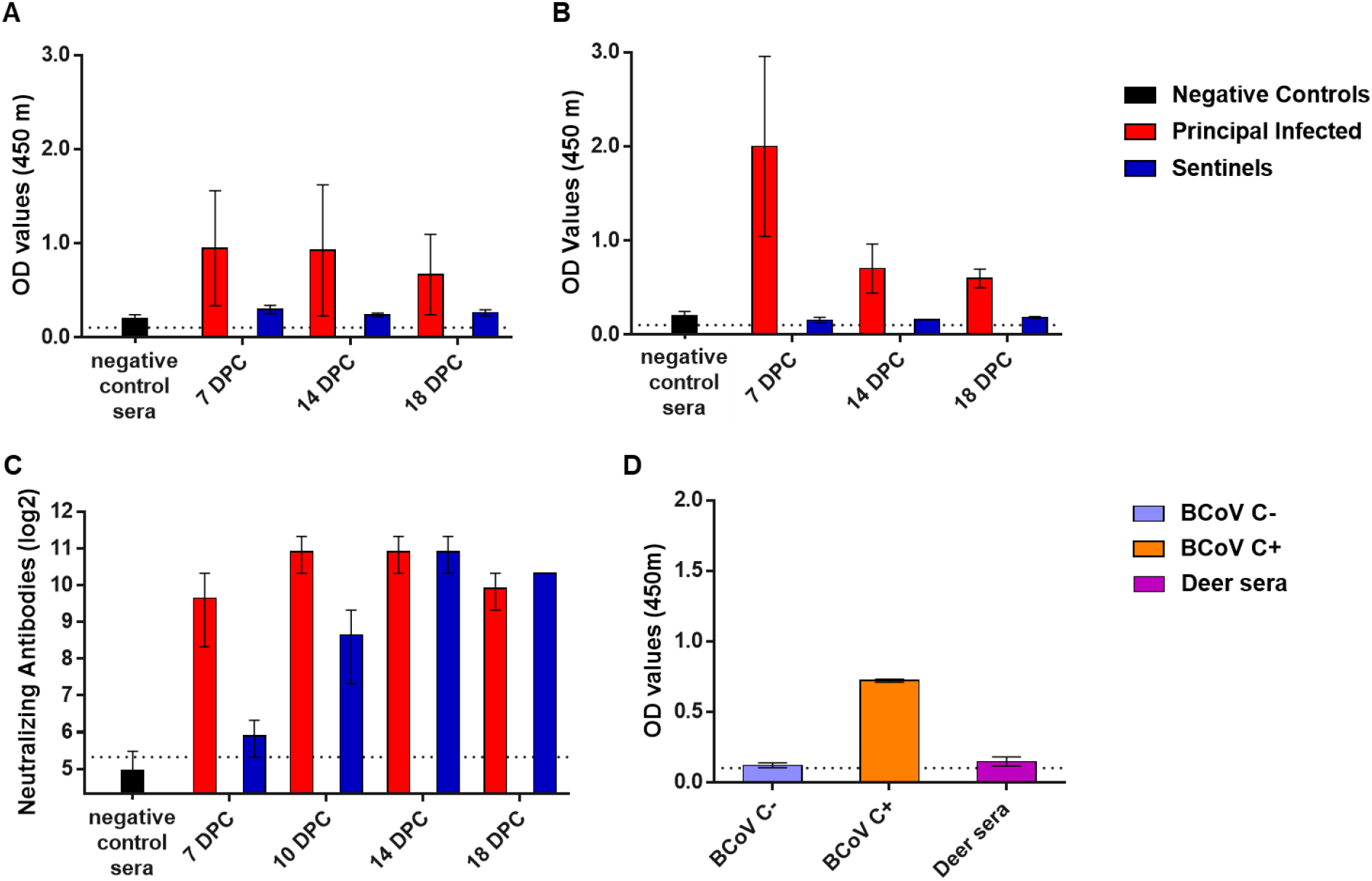
Serology of SARS-CoV-2 infected deer. Detection of SARS-CoV-2 nucleocapsid protein (**A**), and the receptor binding domain (**B**) by indirect ELISA tests. The cut-off was determined by averaging the OD of negative serum + 3X the standard deviation as indicated by the dotted line. All samples with resulting OD values above this cut-off were considered positive. (**C**) Virus neutralizing antibodies detected in serum are shown as log2 of the reciprocal of the neutralization serum dilution. The cut-off of 1:40 is indicated by the dotted line. **A-C**: Negative controls were deer sera (n=4) collected from deer enrolled in a previous epizootic hemorrhagic disease virus vaccine study (36) and pre-challenge sera (−3 DPC) from the six deer enrolled in this study. (**D**) Sera from principal infected (n=4) and sentinel deer (n=2) were tested against the bovine coronavirus (BCoV) spike protein using an indirect ELISA; both, positive (C+) and negative (C-) bovine control sera were included. The cut-off was determined by averaging the OD of negative serum + 3X the standard deviation as indicated by the dotted line. **A-D:** Mean with standard error are shown.

### Competition between SARS-CoV-2 strains in co-infected white-tailed deer

To evaluate the *in vivo* competition between the ancestral lineage A (SARS-CoV-2/human/USA/WA1/2020) and the alpha VOC (SARS-CoV-2/human/USA/CA-5574/2020) strains, cDNA products of SARS-CoV-2 RNA extracted from swabs and tissue homogenates were sequenced on the Illumina NextSeq platform. Sequencing analysis showed the inoculum used for infection was 60% alpha VOC B.1.1.7 and 40% lineage A WA1. Nasal swabs collected at 1 DPC revealed the proportion of lineage A strain ranged from 0.7-40.0% and of alpha VOC B.1.17 strain from 60.0-99.3% in the 4 principal infected deer (**Table 4, Supplementary Figure 4**). In contrast, one of the oral swabs from a principal infected deer (#905) at 1 DPC showed 51.0% lineage A WA1 and 49.0% VOC B.1.1.7, while the other swab contained 5.4% lineage A and 94.6% VOC B.1.1.7. By 3 DPC, the composition of all nasal and most oral swabs was entirely alpha VOC B.1.1.7, with the exception of a 4 DPC oral swab collected from deer #905 that showed 2.9% lineage A and 97.1% VOC B.1.1.7. Swab and tissue samples collected from sentinel deer were also dominated by the alpha VOC B.1.1.7 strain. The B.1.1.7 strain was also dominant in tissues collected from the two primary challenged deer at 4 DPC, with the highest lineage A percentage (20.9%) found in the nasopharynx of deer #905 (**Table 4, Supplementary Figure 4**). Virus genome analysis was only possible in a few tissues collected from principal (nasopharynx and tonsil) and sentinel (retropharyngeal lymph node) deer at 18 DPC; the results revealed a > 95% presence of alpha VOC B.1.1.7 genomic sequences. These results indicate a competitive advantage of the alpha VOC B.1.1.7 isolate over the lineage A WA1 isolate.

**Table 4.**
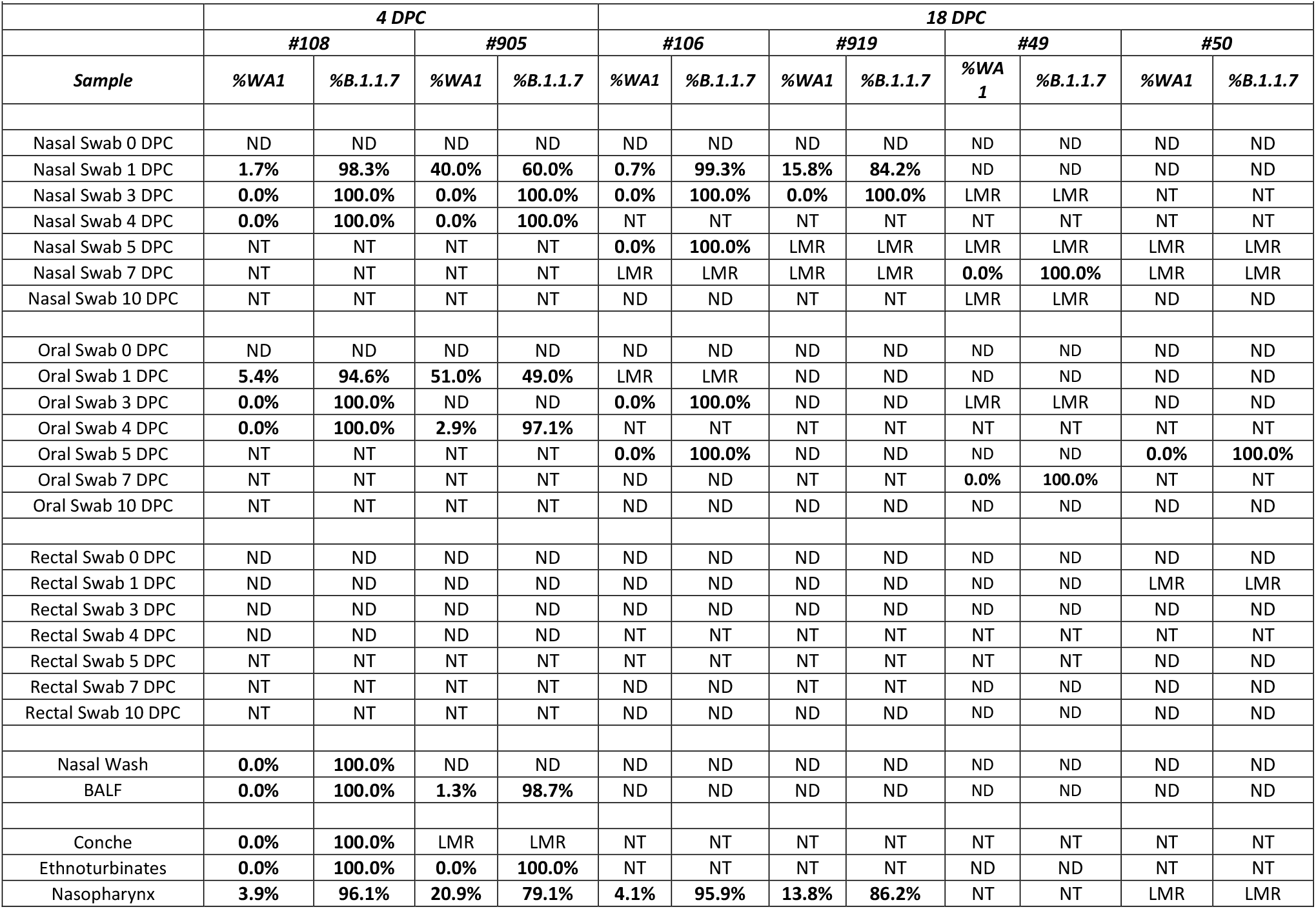
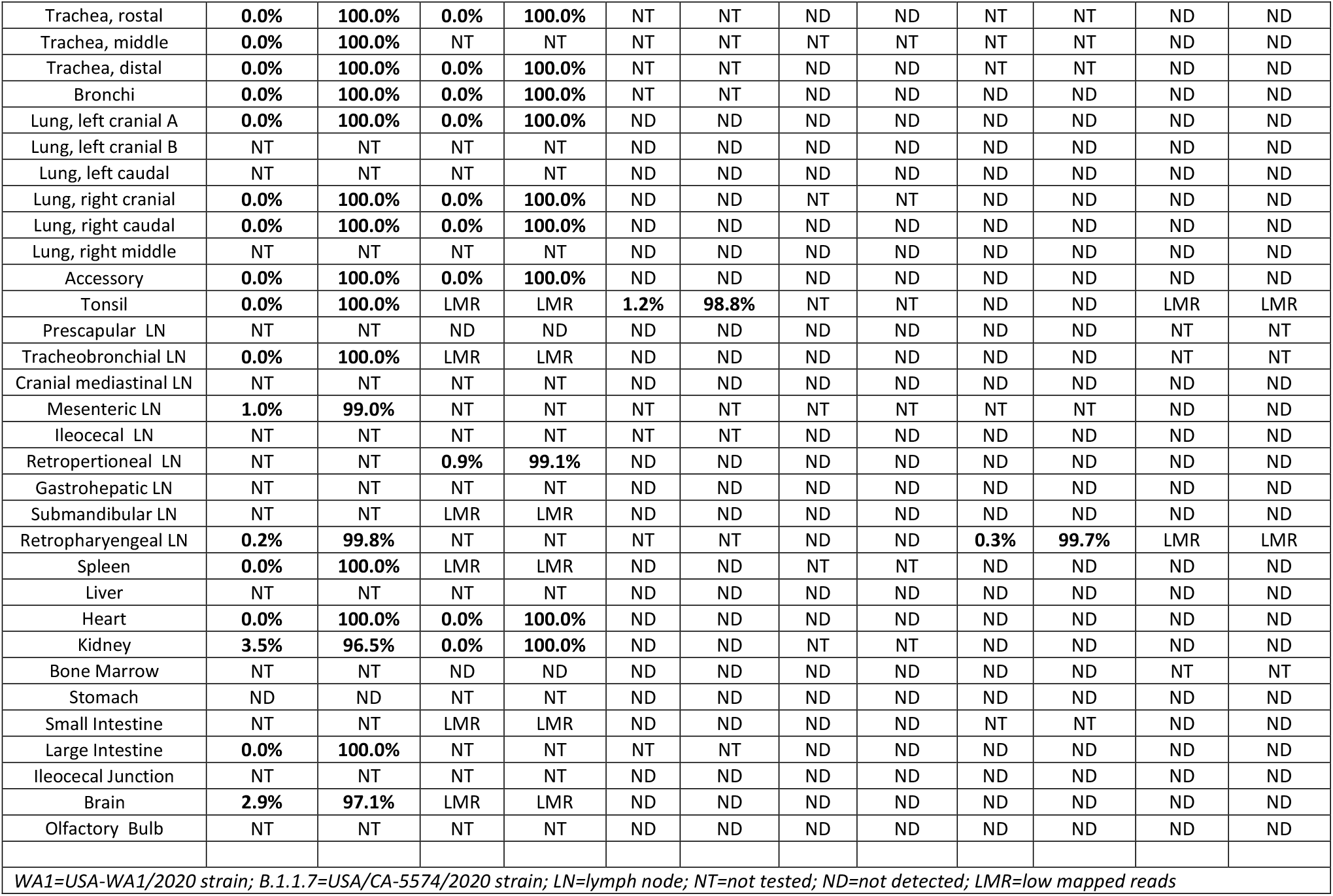
SARS-CoV-2 competition in deer co-infected with lineage A and B.1.1.7 strains determined by next generation sequencing.

## Discussion

Zoonotic diseases reportedly account for approximately 60% of emerging infectious disease (EID) events over the last century^13^. Increased interaction with wild animal populations is a critical factor associated with increased occurrence of zoonotic EIDs. In the last twenty years, three virus species from the genus *Betacoronavirus*, family *Coronaviridae*, have spilled over from wild animal populations into humans, resulting in outbreaks of respiratory disease and global economic losses^4,14,15^. The emergence, global dissemination, and rapid evolution of SARS-CoV-2 has had global impacts on public health and economic stability. The emergence of variant strains, in part due to the exceptional spread of the virus and secondary zoonotic transmissions, has caused additional waves of SARS-CoV-2 infection and reinfection, highlighting the need for enhanced mitigation strategies^3^. To design more comprehensive surveillance efforts, it is necessary to identify susceptible animal hosts^4^ which may contribute as secondary reservoirs to the continued animal-animal and animal-human transmission of SARS-CoV-2. Maintenance of SARS-CoV-2 in wild animal populations would create additional challenges in surveillance and control of this virus. Human interaction with infected wild animal populations could provide avenues for un-checked spillover back into human populations and spontaneous resurgence of SARS-CoV-2 infections with new animal-adapted SARS-CoV-2 variants, as already experienced with mink^3^.

White-tailed deer (*Odocoileus virginianus*) are the second most abundant, densely populated, and geographically widespread wild ruminant species in the United States. Furthermore, commercially farmed deer is a growing industry in the US. Human interaction with white-tailed deer have resulted in the occurrence of infectious diseases such as brucellosis or tuberculosis in human populations in the past^16^. Recently, white-tailed deer fawns were demonstrated to be susceptible to experimental SARS-CoV-2 infection^6^. Also, it was recently reported that sero-surveillance studies conducted in 2021 show that 40% of wild white-tailed deer tested across the midwestern US were positive for SARS-CoV-2 neutralizing antibodies^17^. Although it is unclear at this time if these results are caused by SARS-CoV-2 exposure or cross-reactivity of a yet unidentified closely related coronavirus infection in deer^17^. Surprisingly, antibodies were detected in one sample as early as 2019, and several from January to March of 2020 in that survey, but not in sera collected in years prior to 2019^17^. In the present study, we investigated the susceptibility of approximately 2-year-old, adult white-tailed deer to SARS-CoV-2 infection and the potential for direct transmission to naïve contact sentinel deer. Furthermore, our data is the first to date to provide evidence of vertical transmission of SARS-CoV-2 from mother to fetus. In addition, this study examined the competition of two SARS-CoV-2 isolates, representatives of the ancestral lineage A (SARS-CoV-2/human/USA/WA1/2020; lineage A WA1) and the alpha VOC B.1.1.7 (SARS-CoV-2/human/USA/CA-5574/2020; VOC B.1.1.7) by co-infection of white-tailed deer. We also demonstrated *in vitro* virus replication in primary lung cells derived from white-tailed deer, mule deer and elk, which suggests that besides white-tailed deer, mule deer may also be susceptible to SARS-CoV-2. Elk primary lung cells were refractory to SARS-CoV-2 infection with the ancestral lineage A WA1 strain.

The recently published study by Palmer and coworkers^6^ showed infection and transmission of the SARS-CoV-2 tiger isolate TGR/NY/20 at a dose of 5 × 10^6.3^ TCID_50_ in 6-week-old fawns, with prolonged viral RNA shedding detected in nasal swabs over the course of the 21-day study. Here, we used a lower infectious dose (1 × 10^6^ TCID_50_) in 2-year-old adult deer. Our data indicate that adult deer shed SARS-CoV-2 virus and RNA for a lesser period of time, which could be due to difference in challenge dose, differences in virus isolates, route of administration (IN and PO vs. IN only) or the effects of a more rapid immune response in adult deer. Both studies show lower levels of intermittent virus/RNA shedding detected from oral and rectal swabs through 7 DPC, compared to nasal swabs which were RNA positive for all animals up to 10 DPC in our study. Similarly, as described by Palmer and coworkers^6^ in fawns, we observed efficient transmission of virus to co-housed naïve adult deer, as demonstrated by isolation of SARS-CoV-2 virus from contact sentinels on days 4- and 6-days post comingling (i.e., 5 and 7 DPC) as well as consistent viral RNA detection in swabs and tissues.

In the present study, high copy numbers of viral RNA were detected in many clinical samples and tissues derived from the principal infected and sentinel deer, however virus isolation was infrequent. Infection with the alpha VOC B.1.1.7 has been associated with an abundant production of the nucleocapsid N protein, creating prominent levels of N-specific mRNA which could explain why an N-gene targeting RT-qPCR assay (CDC N2 RT-qPCR assay) might over-estimate viable virus quantities^18,19^. Weak reactivity was observed with some of the pre-challenged deer sera by our *in-house* SARS-CoV-2 N and RBD ELISAs, as well as using an ELISA specific for the bovine coronavirus S protein. Also, some of the deer in our study had borderline levels of SARS-CoV-2 neutralizing antibodies present at pre-challenge. Since these were low levels of antibodies, and all the deer in this study were highly susceptible to SARS-CoV-2 infection, we consider it unlikely that the animals were previously infected with SARS-CoV-2. Nevertheless, we cannot exclude the possibility that the deer were exposed to an unknown deer *Betacoronavirus* with some level of antigenic cross reactivity to SARS-CoV-2, since deer coronaviruses similar to bovine coronaviruses have been previously described^20^; further investigation would be required to ascertain whether this is the case.

Tissues which were collected *postmortem* at 4 DPC and 18 DPC demonstrate widespread distribution of SARS-CoV-2 RNA in the upper respiratory and lymphatic tissues. To evaluate the acute stage of infection, two deer - one pregnant, one not pregnant - were sacrificed at 4 DPC. There was a pronounced difference of viral RNA load (copy number [CN]/mg) in respiratory tissues between these two deer, which may represent different levels of susceptibility between pregnant (low viral loads) and non-pregnant (high viral loads) animals.

Histological evaluations in the upper and lower respiratory tract tissues collected at 4 DPC revealed pathological changes described as rhinitis, marked attenuation of the respiratory epithelium of the trachea, bronchitis, and in some cases bronchiolitis. No interstitial pneumonia was observed. IHC analyses for the presence of SARS-CoV-2 antigen in the respiratory tract at 4 DPC revealed that viral antigen was not detected in the nasal passages, but in the trachea, the bronchi and occasionally in terminal bronchioles. It is suspected that the detection of antigen in only one (#108) of two animals may be due to genetic differences between the respective deer or a susceptibility characteristic of pregnant (#905) versus non-pregnant ($108) deer. In contrast, the histologic examination of principal infected deer euthanized at 18 DPC showed minimal changes in the lungs with no viral antigen detected along the respiratory tract; in the case of sentinel animals, no viral antigen was associated with the histological changes present in the trachea. These findings are not unexpected since none of the deer had obvious clinical signs of disease. As shown previously, cats experimentally infected with SARS-CoV-2 displayed mild to moderate histopathological changes (rhinitis, tracheal and bronchial adenitis) accompanied by viral RNA and antigen at 4 and 7 DPC. However, by 21 DPC histological changes were unremarkable and no detectable viral RNA or antigen was observed^11^.

Sequence analysis preformed on the NextSeq Illumina platform revealed a competitive advantage of the alpha VOC B.1.1.7 (SARS-CoV-2/human/USA/CA-5574/2020) isolate over the lineage A WA1 (SARS-CoV-2/human/USA/WA1/2020) isolate. This data confirms what others have recently reported *in vivo* in ferrets, hamsters, and transgenic ACE2 mice models^5,21^. It is suggested that a component of this competitive advantage is associated with an Asp→Gly substitution at amino acid position 614 (D614G), which alters the interaction of the spike (S) glycoprotein with the hACE2 receptor^22^. The D614G substitution has been shown to enhance the affinity of S to bind hACE2 when compared to the parental D614 strain^5^. Moreover, the N501Y substitution and perhaps even some other S substitutions, might further increase affinity for the ACE2 receptor^23,24^. Additionally, recent data suggests that infection with B.1.1.7 antagonizes innate immunity early in infection by upregulating gene segments which effectively decrease host IFNβ expression and secretion^18^. This allows for uninterrupted viral replication until subsequent changes in transcriptional regulation of SARS-CoV-2 reverse this effect, inducing an inflammatory response which leads to the presentation of typical clinical symptoms and at the same time promotes virus transmission. This sequence of events may partially explain why alpha VOC B.1.1.7 demonstrates higher replication and transmission efficiency than lineage A^18^.

Animal coronaviruses are not commonly associated with reproductive problems, except avian coronaviruses^25,26^. A feline alphacoronavirus which is responsible for feline infectious peritonitis, can be transmitted vertically with major consequences on post-partum kittens resulting in a mortality up to 100% following birth of infected kittens^27^. In the present study, we identified SARS-CoV-2 genetic markers in two fetuses of one principal infected deer euthanized at 4 DPC as evidenced by viral RNA detected from multiple fetal tissues. Interestingly, we only detected viral RNA in the uterus of the deer which was not pregnant, but also had higher viral RNA levels detected systemically in tissues compared to the pregnant doe necropsied at the same time point 4 DPC. We did not detect viral RNA in fetal tissues derived from nine fetuses of does necropsied at 18 DPC, and more than 50% (5/9) of the fetuses collected at this time point were found to be unviable. However, it is difficult to draw conclusively the full effects of SARS-CoV-2 infection in pregnant deer given the small number of animals, the short duration of this study and the absence of control pregnant non-infected animals.

It is well known that pregnancy can increase the risk of illness caused by viral infections like influenza as well as enhance complications due to other underlying medical conditions such as diabetes. The effects of SARS-CoV-2 infection on pregnant women, the pregnancy, the fetus, and *postpartum* are still not fully understood. Several retrospective studies have shown pregnancy to be associated with higher risk of severe illness caused by COVID-19 as well as increased admissions to intensive care and neonatal units; also increased numbers of preterm births (under 37 weeks) have been observed during the course of the pandemic^28–32^. Still births and neonatal death appear to be relatively low for mothers independent whether they are infected or not with SARS-CoV-2^29^. Importantly, in utero SARS-CoV-2 vertical transmission from mother to fetus, while rare, seems possible^33,34^.

Recent large-scale SARS-CoV-2 surveillance efforts in animal and humans have found evidence of reverse-zoonosis (human-animal) resulting in natural infections in companion animals, farmed mink, primates and large cat species in several countries. The source of infection in these incidences are likely infected pet-owners and animal care workers on farms and at zoos. Additionally, SARS-CoV-2 has demonstrated its ability to rapidly mutate and cross back into human populations, giving rise to new variants of interest or concern^3,35^. These situations exemplify the critical need to evaluate potential host species for SARS-CoV-2 which may act as reservoirs for future, secondary zoonotic events. Furthermore, this work demonstrates the need for more intensive, focused surveillance efforts on high-risk animal populations, such as farmed and wild white-tailed deer and mule deer populations, as well as farm, wildlife and zoo workers, in order to identify new animal derived SARS-CoV-2 variants which may evade current mitigation strategies.

## Acknowledgments

We thank the staff of KSU Biosecurity Research Institute, the histology laboratory at the Kansas State Veterinary Diagnostic Laboratory (KSVDL), members of the Histology and Immunohistochemistry sections at the Louisiana Animal Disease Diagnostic Laboratory (LADDL), the Comparative Medicine Group staff at Kansas State University and technical support from Emily Gilbert-Esparza, Yonghai Li, and Jingwen Peng of KSU, and Dane Jasperson from USDA. The SARS-CoV-2 strains USA/CA-5574/2020 and USA/WA1/2020 were obtained through BEI Resources (catalog # NR-52281 and #54011). We also thank Dr. Kyeong-Ok Chang for the Vero E6/TMPRSS2 cells used in these studies.

## SUPPLEMENTARY FIGURES 1-4

**Supplementary Figure 1.**
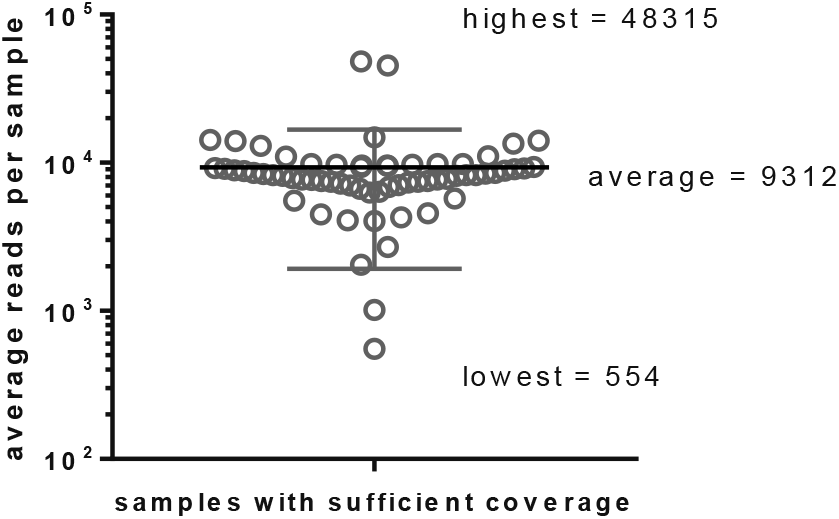
Read coverage of sequenced samples from SARS-CoV-2 co-infected white-tailed deer. Swab and tissue homogenate samples from white-tailed deer co-infected with the SARS-CoV-2/human/USA/WA1/2020 (lineage A WA1) and SARS-CoV-2/human/USA/CA-5574/2020 (alpha VOC B.1.1.7) strains were analyzed using next generation sequencing.

**Supplementary Figure 2.**
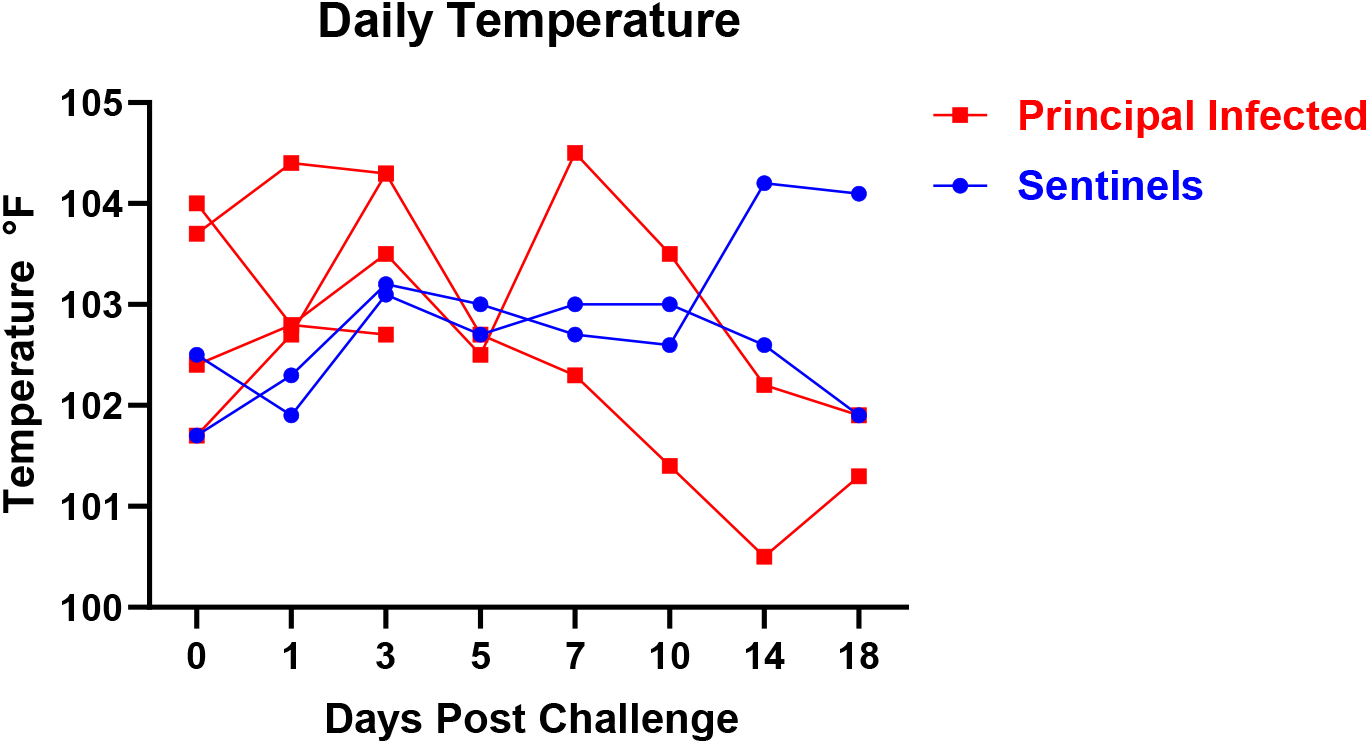
Daily Temperatures. Rectal temperatures were taken from sedated deer on 0, 1, 3, 5, 7, 10, 14, 18 DPC.

**Supplementary Figure 3.**
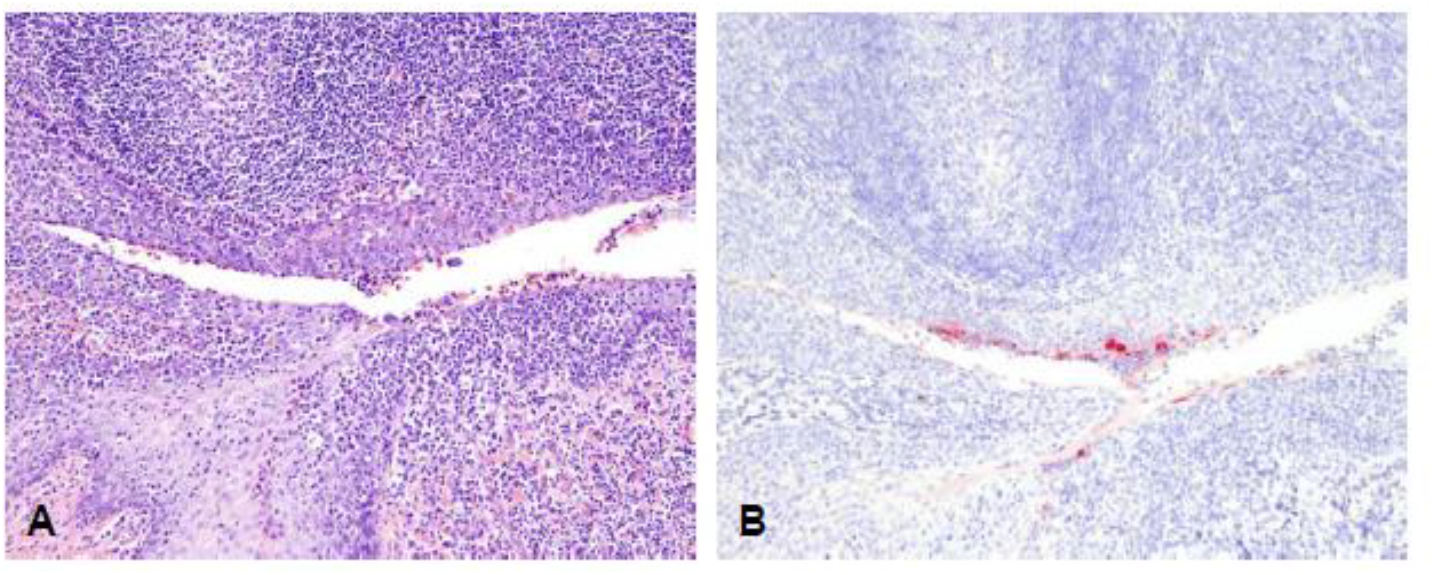
Histopathological lesions in the tonsil of a primary infected deer at 4 DPC. Histologic alterations and SARS-CoV-2 antigen distribution in the tonsil of a white-tailed deer at 4 DPC. Foci along the lining epithelium are characterized by erosion and sloughing of superficial layers of the stratified squamous epithelium, with moderate numbers of transmigrating neutrophils and lymphocytes (**A**), and few superficial epithelial cells containing viral antigen (**B**). H&E and Fast Red, 100X total magnification.

**Supplementary Figure 4.**
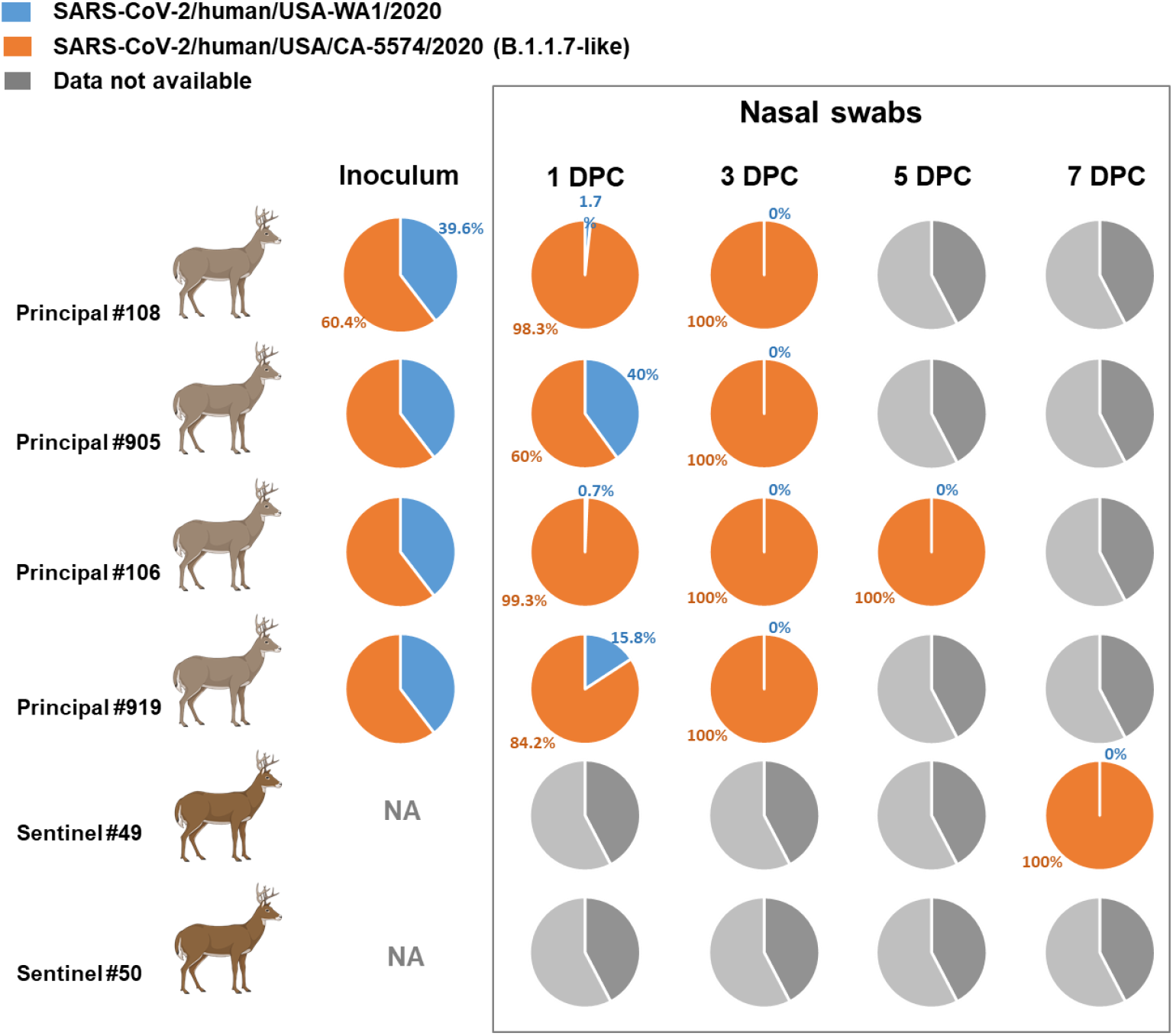
Next-generation sequencing of swabs collected from SARS-CoV-2 co-infected white-tailed deer. cDNA products of SARS-CoV-2 RNA extracted from nasal swabs were sequenced on the Illumina NextSeq platform to evaluate the *in vivo* competition between the ancestral lineage A WA1 (SARS-CoV-2/human/USA/WA1/2020) and the alpha VOC B.1.1.7 (SARS-CoV-2/human/ USA/CA-5574/ 2020) strains.

